# A high throughput platform for measuring and predicting vitrification behavior in multicomponent aqueous solutions

**DOI:** 10.64898/2026.02.19.706831

**Authors:** Nima Ahmadkhani, Cameron Sugden, Euihyun Lee, Carlos R. Baiz, Duncan Brown, Nolan Drummond, Alec Snyder, Matthew Uden, Adam Z. Higgins

**Author notes:** N.A. and C.S. contributed equally to this work.

## Abstract

Cryopreservation depends critically on suppression of ice formation by cryoprotective agents (CPAs), but limited data is available on the CPA concentration required for vitrification (Cv). Here, we introduce a high-throughput 384-well platform that integrates automated liquid handling, randomized plate layouts, and a binary-search strategy to rapidly determine Cv across hundreds of formulations. Relative to conventional methods, this approach increases throughput by ∼50-fold, compressing a year of measurements into one week, while markedly reducing manual labor. Across ∼200 CPA compositions, we demonstrate that environmental boundary conditions strongly influence vitrification behavior: plates sealed with silicone mats exhibited lower Cv than open plates, indicating that sealed configurations promote vitrification. Further, the data reveal a decrease in Cv with increasing CPA molecular weight, consistent with enhanced ice suppression by larger molecules. We also present a simple mixture model that accurately predicts Cv for a broad range of CPA formulations, including mixtures containing up to seven CPAs (R² ≥ 0.93), and use this model to evaluate published CPA toxicity data to identify formulations that operate near their vitrification threshold while maintaining relatively low toxicity. Together, these results establish a framework for rapid Cv determination, predictive modeling of vitrification behavior, and rational design of CPA formulations.

## 1 Introduction

Interest in cryopreservation of complex systems has grown substantially over the past decade, driven by its potential impacts across fields ranging from organ transplantation^1, 2^ to conservation of endangered species^3–5^ to suspended animation for long-term space travel^2, 6^. This momentum has been fueled by recent reports of successful cryopreservation of rat and rabbit kidneys with functional recovery after transplantation^7–10^, and the recent demonstration of vitrification without ice formation or cracking at human organ scale^11^.

While advances in rapid warming technologies have contributed to recent successes with small animal organs, human organs face a bottleneck during cooling (Figure 1A). Their larger size leads to slower cooling, increasing the risk of ice formation. Preventing ice under these conditions requires higher concentrations of cryoprotective agents (CPAs), which in turn elevates the risk of toxicity. This tradeoff between toxicity and stability against ice formation is the central challenge of vitrification-based cryopreservation.

**Figure 1.**
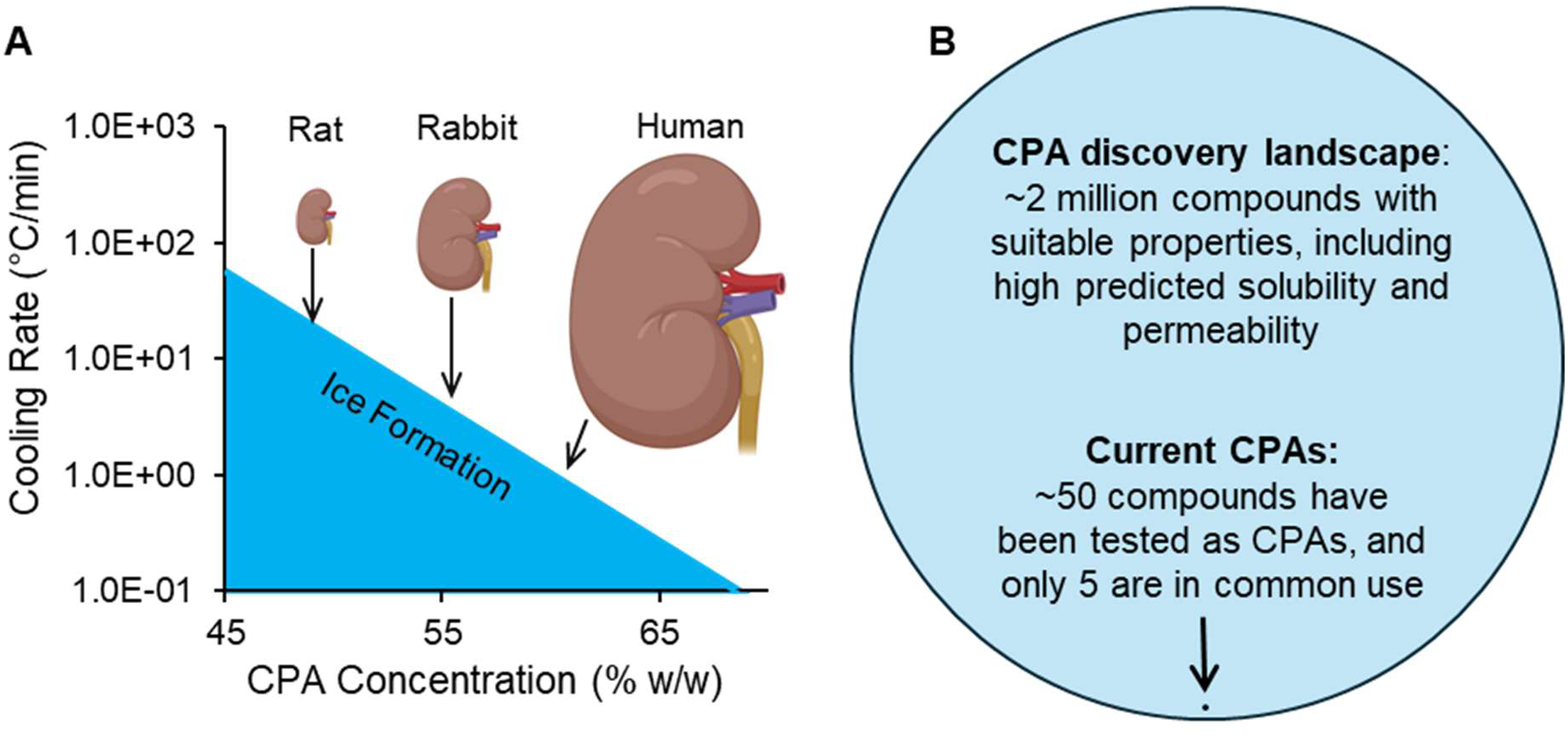
Challenges and opportunities for cryopreservation of complex systems. (A) Despite recent successes with cryopreservation of rat and rabbit kidneys, challenges remain with translating this to human organs due to their larger size and slower achievable cooling rate. To prevent ice formation at slower cooling rates, higher CPA concentrations are needed, which increases the risk of toxicity. This underscores the need for new CPA formulations that are stable against ice formation and nontoxic. Ice formation boundary predicted as described previously.^14^ (B) Current CPAs only scratch the surface of the CPA discovery landscape, highlighting the huge opportunity to find new and improved CPA formulations. Of the >100 million compounds in the PubChem database, we estimate that ∼2 million have potential as CPAs. Yet only ∼50 compounds have been tested as CPAs and only 5 are in common use.

To overcome this challenge, there is tremendous interest in developing new CPA formulations that provide better vitrification stability with less toxicity. Contemporary vitrification protocols typically rely on a small set of CPAs, such as dimethyl sulfoxide, formamide, ethylene glycol, glycerol, and propylene glycol, often used in mixtures that balance stability, viscosity, and toxicity.^7, 8^ These compounds represent only a narrow portion of the accessible chemical space (Figure 1B). Public compound resources contain hundreds of millions of small molecules,^12, 13^ and we estimate that there are millions with suitable properties for use as CPAs. Many of these molecules, or their mixtures, could improve stability against ice formation while also reducing toxicity, yet they remain almost entirely unexplored.

To explore this large chemical space, high throughput methods are needed to assess key properties including toxicity, cell membrane permeability, and vitrification stability. Recent work has introduced high-throughput platforms for screening CPA toxicity and cell membrane permeability.^15–19^ However, no analogous high-throughput method exists for evaluating stability against ice formation. This is a major bottleneck for large-scale screening to discover new CPAs and optimize CPA formulation.

A common parameter for characterizing stability is the minimum vitrifying concentration (Cv). In simple terms, Cv is the lowest CPA concentration that prevents any detectable ice when the sample is cooled along a defined temperature profile and set of environmental conditions. The most common approach for measuring Cv involves plunging individual tubes of CPA solution into liquid nitrogen and assessing ice formation visually^20^ or via X-ray diffraction.^21^ This approach requires multiple manual steps (filling tubes, sealing, plunging, and post-plunge inspection), and each tube represents a single data point. Consequently, the overall throughput is low, which helps explain why the largest published Cv dataset is limited to 45 CPA solutions.^22^

To address this gap, we developed a high-throughput method that enables quantification of Cv in a 384-well format under cooling conditions consistent with human organ cryopreservation. This new approach increases throughput by ∼50x and significantly reduces manual labor via automation of liquid handling and data analysis. We use this new approach to generate ∼400 Cv measurements based on ∼26,000 individual data points, enabling examination of molecular and environmental factors that influence vitrification behavior. In addition, we present a model for predicting Cv in CPA mixtures, and use the model to examine published toxicity data to identify CPA formulations with low toxicity in the concentration regime near Cv.

## 2 Materials and methods

### 2.1 Materials

We used 14 CPAs in this study (Table 1), corresponding to the same set of chemicals previously screened by Ahmadkhani et al.^15^ for CPA toxicity. These CPAs were chosen because they are cell membrane permeable and have relatively low toxicity. All CPA solutions were prepared in deionized water supplemented with 3.2% w/v dextrose (D-glucose) (Fisher Chemical).

**Table 1.** List of CPAs and corresponding abbreviations.

| No | CPA Name (Abbreviation) | Vendor | Purity |
| --- | --- | --- | --- |
| 1 | 1,3-Dihydroxyacetone (DHA) | Ambeed | 98% |
| 2 | 1,3-Propanediol (PD) | TCI America | 98% |
| 3 | 2,3-Butanediol (BD)* | MilliporeSigma | 98 % |
| 4 | 2-Methoxyethanol (ME) | Thermo Scientific Chemicals | ≥99% |
| 5 | 2-Methyl-1,3-Propanediol (MP) | TCI America | ≥98% |
| 6 | Acetamide (AM) | Thermo Scientific Chemicals | 99% |
| 7 | Diethylene Glycol (DG) | VWR | 99% |
| 8 | Dimethyl Sulfoxide (DMSO) | Thermo Fisher Scientific | 99.7% |
| 9 | N,N-Dimethylacetamide (DMA) | Sigma Aldrich | ≥99% |
| 10 | Ethylene Glycol (EG) | Macron Fine Chemicals | ≥99% |
| 11 | Formamide (FA) | VWR | 99 % |
| 12 | Glycerol (Gly) | Macron Fine Chemicals | 99.5% |
| 13 | N-Methylacetamide (NMA) | TCI America | ≥99% |
| 14 | Propylene Glycol (PG) | VWR | 99.5 % |

\* Mixture of meso-, D-, and L-forms

### 2.2 Preparation of CPA stock solutions

CPA stock solutions (70% w/v) were prepared by weighing 35 ± 0.05 g of the pure CPA into a 50 mL conical tube, adding 5 mL of 32% w/v dextrose stock, and bringing the final volume to 50 mL with deionized water. Solutions were vortexed until completely homogenized, resulting in CPA stock solutions at 70% w/v containing 3.2% w/v dextrose.

### 2.3 Automated CPA mixture preparation using robotic liquid handling

CPA mixtures were prepared using a Hamilton Microlab STAR liquid-handling system. CPA stock solutions and CPA-free 3.2% w/v dextrose were first manually transferred into glass tubes. These tubes were then used as the source for automated preparation of CPA mixtures in a 96 deep-well plate (working volume: 1000 µL per well). A custom MATLAB script was used to calculate the required volume of each source tube solution to make the desired CPA mixture, and to write the corresponding commands for liquid transfers in the liquid handler.

The resulting 96-well plate was then used as the source for automated preparation of three 384-well plates. A 70 µL volume of each CPA formulation was dispensed into four randomized wells in each 384-well plate, resulting in 12 total replicates per condition distributed across all three plates. Well locations were randomized to minimize positional bias. As a visual control to verify correct dispensing locations, food dye was included in 4–8 replicate wells per plate. In rare cases, we noted a pipetting error that prevented liquid transfer into some of the wells. This affected 118 wells out of ∼26,000 total (< 0.5%), resulting in a slightly reduced replicate count in some cases (the lowest replicate count was 8). Plates were centrifuged prior to cooling to remove trapped air bubbles.

### 2.4 Cryogenic cooling apparatus for multi-well plates

To enable controlled cryogenic cooling of multi-well plates, a custom cooling platform was constructed (Figure S1). The cooling platform was housed inside a large Styrofoam chamber to reduce environmental heat exchange. The primary cooling element consisted of four aluminum blocks, each with a thickness of 1.125 inches, a width of 6 inches, and a length of 6 inches. The blocks were tiled to create a 1 ft x 1 ft cooling surface. The aluminum blocks were supported by twenty aluminum cylinders (1 inch diameter, 1 inch thick), allowing liquid nitrogen to circulate beneath the blocks. Upon addition of liquid nitrogen, the block surfaces equilibrated near −190 °C. Block temperatures were recorded prior to sample loading.

To promote uniform thermal distribution across the 384-well plate, a multilayer cooling assembly was implemented. This was housed within a stainless-steel steam pan (24 gauge, 2/3 size, 6 inches deep) with a base of ∼1 ft x 1ft that matched the size of the aluminum cooling platform. The three 384-well plates were placed within this pan, with a layer of acrylic and copper below each plate. A 0.476-cm thick acrylic sheet was included as an insulating layer to produce cooling rates representative of those encountered in human organ-scale samples. A custom-cut copper plate, sized to fully contact the underside of the 384-well plate, was positioned above the acrylic sheet to enhance lateral heat spreading and minimize temperature gradients across wells.

To minimize frost formation during cooling, a cube-shaped acrylic cover was placed over the top of the stainless-steel pan, and silica gel desiccant was added at the base of the pan. The interface between the acrylic cover and the stainless-steel pan was wrapped in plastic film to limit vapor exchange and maintain a semi-airtight environment. Plates were placed inside the enclosure for at least 30 minutes prior to cooling to reduce condensation during cooling.

### 2.5 Environmental boundary conditions above wells

Immediately prior to cooling, plates were subjected to different environmental boundary conditions above the wells to assess their effect on vitrification behavior. Plates were either left uncovered (open-top), purged with argon under open-top conditions, or covered using silicone sealing mats or acrylic lids (0.1875 inches), depending on the experimental configuration

### 2.6 Cooling rate measurement

Cooling rates were measured using a K-type thermocouple connected to an Omega HH502 digital thermometer. Cooling rates were calculated as the slope of the temperature drop between −20 °C and −120 °C (Figure S2).

To characterize cooling behavior within the 384-well plate, thermocouple probes were inserted into wells at three representative locations: A1, A12, and H12. These wells were chosen to represent edge and center locations. Small openings were created in the silicone mat at these positions to allow direct probe access while keeping the remainder of the plate sealed. Each location was tested using three different 384-well plates to confirm reproducibility.

For these measurements, plates contained CPA formulations as defined in Table S1 in randomized well locations, resulting in a mixture of wells that vitrified and wells that froze during cooling. This configuration reflects the heterogeneous thermal and phase conditions encountered during routine Cv screening rather than an artificially uniform state.

### 2.7 Effect of plate position and local neighborhood on Cv measurement

To test whether (i) plate position and/or (ii) the dominant phase state in surrounding wells (ice vs glass) biases vitrification outcomes in the 384-well assay, we performed a controlled validation experiment using two representative CPAs: DMSO and Gly.

For each CPA, solutions spanning 40–70% w/v were prepared in 2% w/v increments. DMSO was dispensed into columns 1 and 11, and Gly into columns 13 and 24, to sample both edge and interior regions of the plate. Cv was estimated separately for each column as the lowest concentration in the column that vitrified.

Two covering configurations were tested: a silicone mat and an acrylic lid. For each cover condition, two background (“neighborhood”) environments were created by filling all remaining wells with either a glass-forming solution (70% w/v DMSO) or an ice-forming solution (40% w/v DMSO).

### 2.8 Validation of Hamilton pipetting

To verify the accuracy of dispensing by the Hamilton Microlab STAR robotic system, all eight independent pipetting channels were evaluated using both 1000 µL and 300 µL tip heads. Two representative CPAs were selected to span a range of physical properties: Gly (70% w/v; high viscosity) and PG (70% w/v; low surface tension). For each combination of CPA, tip size, and channel, the dispensed mass was measured using an analytical balance and compared to manual pipetting of the same target volume. Each condition was tested in triplicate. The ratio of the mean mass dispensed by the robot to that obtained by manual pipetting was 0.994 for Gly and 1.003 for PG, with no statistically significant differences observed, confirming accurate and consistent performance across all channels.

The Hamilton platform features customizable liquid classes that let users tailor pipetting parameters to the physical properties of each liquid. Separate liquid classes were implemented for CPA-free aqueous solutions and CPA solutions. Transfers for CPA-free solutions were performed using standard surface-dispense parameters. In contrast, a CPA liquid class was specifically optimized to account for increased viscosity and altered flow behavior relative to water. For CPA handling, aspiration and dispense flow rates were reduced, settling times were introduced, and controlled air-transport volumes were applied to minimize bubble formation, splashing, and dispense variability during high-throughput operation. Pressure-based liquid level detection was disabled to prevent false triggering in viscous solutions.

During transfer from the 96 deep-well plate to the 384-well plate, an aliquoting strategy was used in which all eight independent channels aspirated sufficient volume to dispense into four target wells, with an additional excess volume. Prior to dispensing into the destination wells, 10 µL was dispensed back into the source well to eliminate trapped air or bubbles in the tips. The remaining volume was then distributed to four randomized locations without returning to the source plate. After dispensing, the residual 10 µL was returned to the source well. This approach ensured bubble-free operation, minimized dead volume, and improved volumetric accuracy and reproducibility during high-throughput transfer.

### 2.9 Binary search strategy for determining Cv

A traditional linear scanning approach (e.g., testing 1% concentration increments from 38% to 70% w/v) would require 33 measurements to identify Cv. To reduce the number of required experiments, we implemented a binary-search strategy^23^ to iteratively narrow the concentration range (Figure S3).

The search begins at the midpoint of the predefined bounds (38–70% w/v), resulting in an initial test concentration of 54% w/v. Each formulation was evaluated at this concentration using 12 replicates, and the next concentration tested was determined based on the fraction of the wells that vitrified using a threshold of 0.75. If the vitrified fraction was less than 0.75, then the next concentration was set to the midpoint between 54% w/v and the upper bound (70% w/v), yielding 62% w/v. If the vitrified fraction was greater than or equal to 0.75, the next concentration tested was the midpoint between 38% and 54% w/v (i.e., 46% w/v). This iterative halving process continued until the concentration at the vitrification boundary was resolved. Using this approach, only five iterations were required to achieve a resolution of 1% w/v, substantially reducing experimental effort compared with linear scanning.

Custom code in Python and MATLAB was used to automate calculation of the new concentrations to test in each iteration, as well as creation of an input file for the Hamilton liquid handler containing commands for preparing the corresponding CPA solutions.

After completing 5 iterations of binary search, the resulting concentration series was fit using a sigmoid model to estimate Cv:

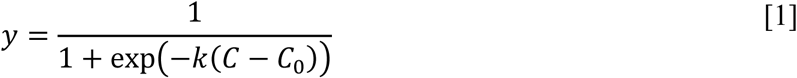

where *y* is the vitrified fraction, *C* is the total CPA concentration, *C*_o_ is the inflection point and *k* governs the steepness of the transition. Least squares fitting was performed using the Levenberg-Marquardt algorithm. Cv was estimated as the concentration at which the best-fit sigmoid model reached the vitrification threshold. We use a threshold of 0.75 here, but the data file in Supporting Information also provides Cv for a threshold of 0.99. We did not fit a sigmoid model for cases where the vitrification fraction was lower than 0.67 over the entire range of concentrations tested.

Uncertainty was quantified using non-parametric bootstrapping with 1000 samples for each CPA mixture formulation. For each bootstrap replicate, the binary observations (vitrified or frozen) at each concentration were randomly resampled with replacement, the vitrified fraction at each concentration was computed, the sigmoid model was fitted, and Cv was calculated using the best-fit sigmoid model. This resulted in a Cv estimate for each of the 1000 bootstrap replicates. The distribution of Cv values was used to compute 95% confidence intervals as the 2.5^th^ and 97.5^th^ percentiles. For sigmoid model fitting we used a Gauss-Newton solver with the following filters applied: (1) bootstrap replicates were only accepted if the sigmoid fit improved the residual sum of squares compared to a baseline model equal to a horizontal line at the mean vitrified fraction, (2) bootstrap replicates were only accepted if Cv was higher than the highest concentration with a vitrified fraction of zero. Overall, only 0.01% of bootstrap replicates were rejected by the filters.

### 2.10 Image analysis

Plate images acquired at the end of cooling were analyzed using a custom Python-based workflow. Raw photographs were first imported in RGB format. To ensure accurate well detection, users interactively indicated the centers of four reference wells (top-left, top-right, bottom-left, bottom-right). These points were used to perform bilinear interpolation to generate perspective-corrected coordinates for all 384 wells (16 × 24 array).

For each well, average RGB intensities were computed within a circular region of interest, and the mean and standard deviation of pixel intensities were extracted. Automated classification of vitrified versus frozen wells was performed by normalizing the blue-channel intensity relative to local background and applying dual thresholds based on normalized intensity and intensity variance to identify wells containing ice. A graphical interface allows users to manually review and correct automated classifications by clicking individual wells, providing quality control for ambiguous cases. Automated image analysis classified vitrified wells with 92% accuracy and frozen wells with 66% accuracy, resulting in an overall classification accuracy of 80%. Classification errors primarily arose from wells containing small ice crystals that were challenging for the algorithm to detect. Figure S4 shows representative images from the different stages of the image analysis process.

### 2.11 Mixture model for estimating Cv in multi-CPA solutions

To model Cv in CPA mixtures, we assume that each CPA independently interacts with water to suppress ice formation. For each CPA, we define an intrinsic ice suppression parameter *α* that represents the CPA concentration that interacts with the threshold amount of water required to fully prevent ice formation under the cooling conditions in our Cv experiments. For single-CPA solutions, *α* = *C*_v_if ice is the only crystalline phase that forms. For CPA mixtures at Cv, the net contribution of all CPAs must reach the threshold for vitrification. We assume that each CPA interacts with water in proportion to its intrinsic ice suppression parameter *α*. For a CPA mixture at its vitrification concentration *C*_v_, this results in:

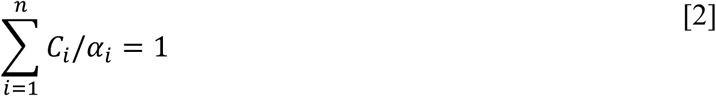

where *C*_i_ is the concentration of the i^th^ CPA and *n* is the number of CPAs in the mixture. The equation sums to 1, which indicates that the CPA mixture is at the threshold for vitrification. To express this in terms of *C*_v_ we can substitute *C*_i_ = *x*_i_*C*_v_ into the above equation, where *x*_i_ is a CPA mole ratio defined as the moles of the i^th^ CPA divided by the total moles of CPA. This results in:

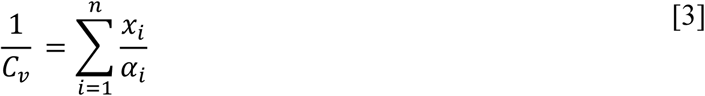

Several single-CPA solutions either did not vitrify over the entire range of concentrations tested or exhibited behavior consistent with alternative crystallization pathways (besides ice). This prevented reliable direct measurement of ice suppression ability based on Cv measurements for single-CPA solutions. To overcome this limitation, we estimated the intrinsic ice suppression parameter *α* for each CPA by fitting Eq. 3 to the data from two-CPA mixtures, where non-ice crystallization is relatively unlikely.

### 2.12 Statistical analysis

Statistical comparisons between experimental groups were performed using a two-tailed Student’s t-test assuming equal variances. For examining relationships between variables, linear regression analysis was used. Analyses were conducted with Microsoft Excel. Statistical significance was defined as p < 0.05.

### 2.13 Molecular dynamics simulations

MD simulations were performed using the GROMACS simulation package^24–26^ employing the CHARMM36 general force field for CPAs^27–30^ with the TIP4P-Ew water model. Simulation procedures are described in a recent article.^31^ In brief, initial configurations of CPA solutions were constructed using the Packmol package.^32^ To investigate the effect of CPA on the water H-bond network structure, 14 single-CPA solutions (EG, AM, PD, ME, PG, GLY, DG, BD, MP, DMSO, DMA, NMA, FA, DHA) were prepared with 4 different concentrations.

Initial configurations of systems were energy-minimized. Following minimization, a 2 ns NVT and 5 ns NPT equilibration were carried out at 298 K. Following equilibration, 10 ns simulated annealing was performed from 298 K to 198 K. Next, 5 ns NVT and 30 ns NPT equilibrations were carried out at 198 K. Final production run was performed for 20 ns under constant NPT conditions. Simulation trajectories were stored every 10 ps for the statistical analysis. The V-rescale thermostat^33^ and the C-rescale barostat^34^ were employed to control simulation temperature and pressure during the entire simulation process. The SHAKE algorithm^35^ was applied.

## 3 Results

### 3.1 High-throughput platform for measuring Cv

To accelerate discovery of CPA formulations suitable for vitrification of complex systems (e.g., organs), we developed a high-throughput Cv screening platform based on an automated 384-well plate workflow (Figure 2). In this system, the Hamilton liquid-handling robot makes the CPA mixtures in a 96-well plate and then dispenses them into randomized well positions in three 384-well plates. The 384-well plates are then placed in a cooling apparatus that enables simultaneous cooling of all three plates. Images acquired at the end of cooling are then subjected to semi-automated analysis to classify the wells as either frozen or vitrified, and the resulting data is fed to a binary search algorithm^23^ for calculating the next set of compositions to test. Compared with conventional tube-based methods, this automated workflow significantly increases Cv measurement throughput, enabling automated preparation of CPA mixtures in three 384-well plates in less than 4 hours, cooling of the well plates in ∼1 h, and semi-automated image analysis in less than 30 min. A single run generates 1,152 individual data points (384 × 3), a scale of measurement that would require weeks to complete using conventional tube-based approaches. Through this integrated approach, we generated approximately 26,000 total data points, corresponding to ∼400 distinct Cv measurements (see Supporting Information for the full dataset). Together, these results provide a comprehensive mapping of vitrification behavior across a broad compositional and concentration space.

**Figure 2.**
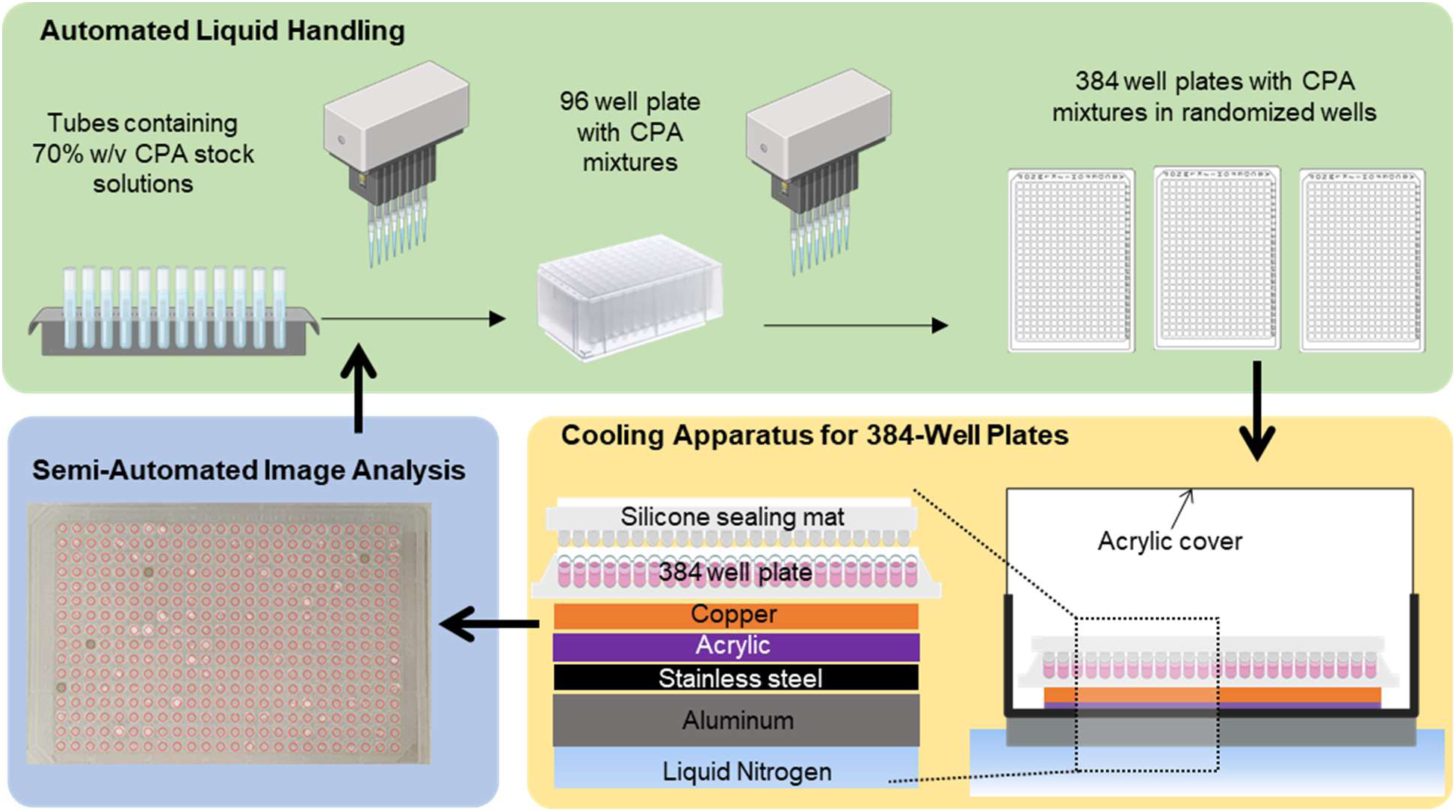
Schematic of the high throughput Cv platform. By combining automated liquid handling for CPA mixture preparation, simultaneous cooling of three 384 well plates, semi-automated image analysis, and binary search^23^ for efficiently homing in on Cv, this setup can achieve a Cv measurement throughput ∼50 times higher than conventional methods.

### 3.2 Cooling-rate characterization across the 384-well plate

A primary design objective of this platform was to measure Cv at cooling rates relevant to organ-scale cryopreservation, rather than at the much higher rates typical of small-volume systems. To achieve this target cooling regime, an insulating acrylic layer was incorporated beneath the 384-well plate to moderate heat transfer and intentionally slow the effective cooling rate experienced by the samples (see Methods).

Cooling rates were quantified at three representative plate locations (A1, A12, and H12) over the temperature range −20 °C to −120 °C. The resulting cooling rates showed modest spatial variation, with values of approximately 2.6 °C/min at A1, 1.6 °C/min at A12, and 1.7 °C/min at H12 (Figure 3; Figure S2). Overall, the average cooling rate was 2.0 ± 0.3 °C/min, which matches the cooling rate achievable in human kidneys ^11^.

**Figure 3.**
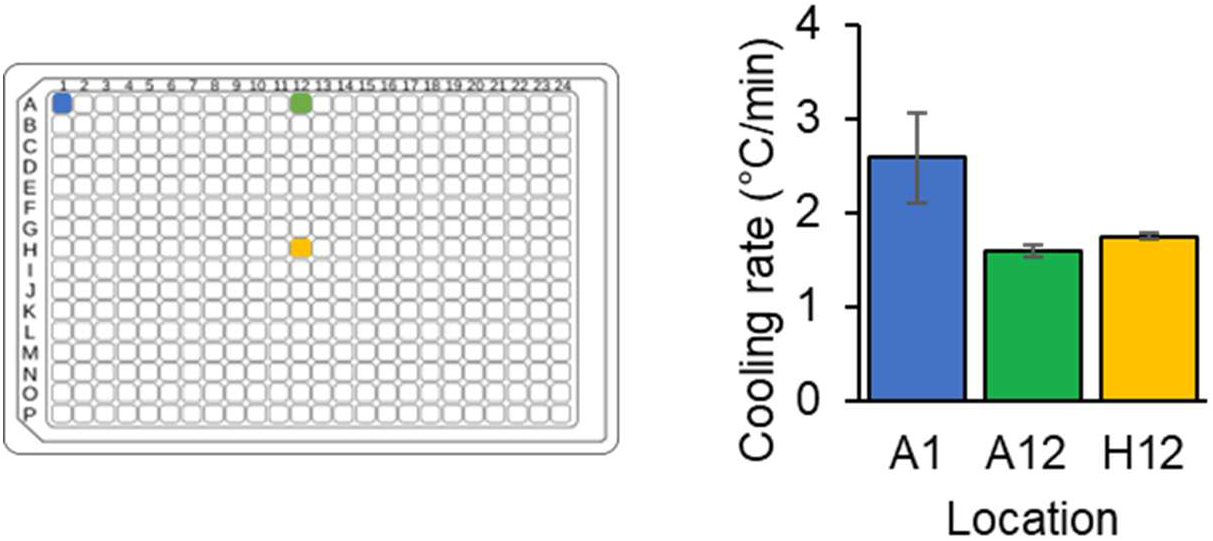
Cooling uniformity in 384-well plates. Cooling rate was measured at three representative well locations (A1, A12, H12), illustrating modest variation in cooling rate across the plate.

### 3.3 Effect of plate position and local neighborhood on Cv measurement

Despite small positional differences in cooling rate, a positional and neighborhood validation experiment revealed no detectable differences in Cv (Figure 4). We measured Cv for DMSO and Gly at central and edge locations under two neighborhood conditions: ice-dominant (with neighboring wells containing 40% w/v DMSO) and glass dominant (with neighboring wells containing 70% w/v DMSO). As shown in Figure 4B and Figure S5, Cv values for both CPAs were indistinguishable as a function of plate position or neighborhood classification (p > 0.05).

**Figure 4.**
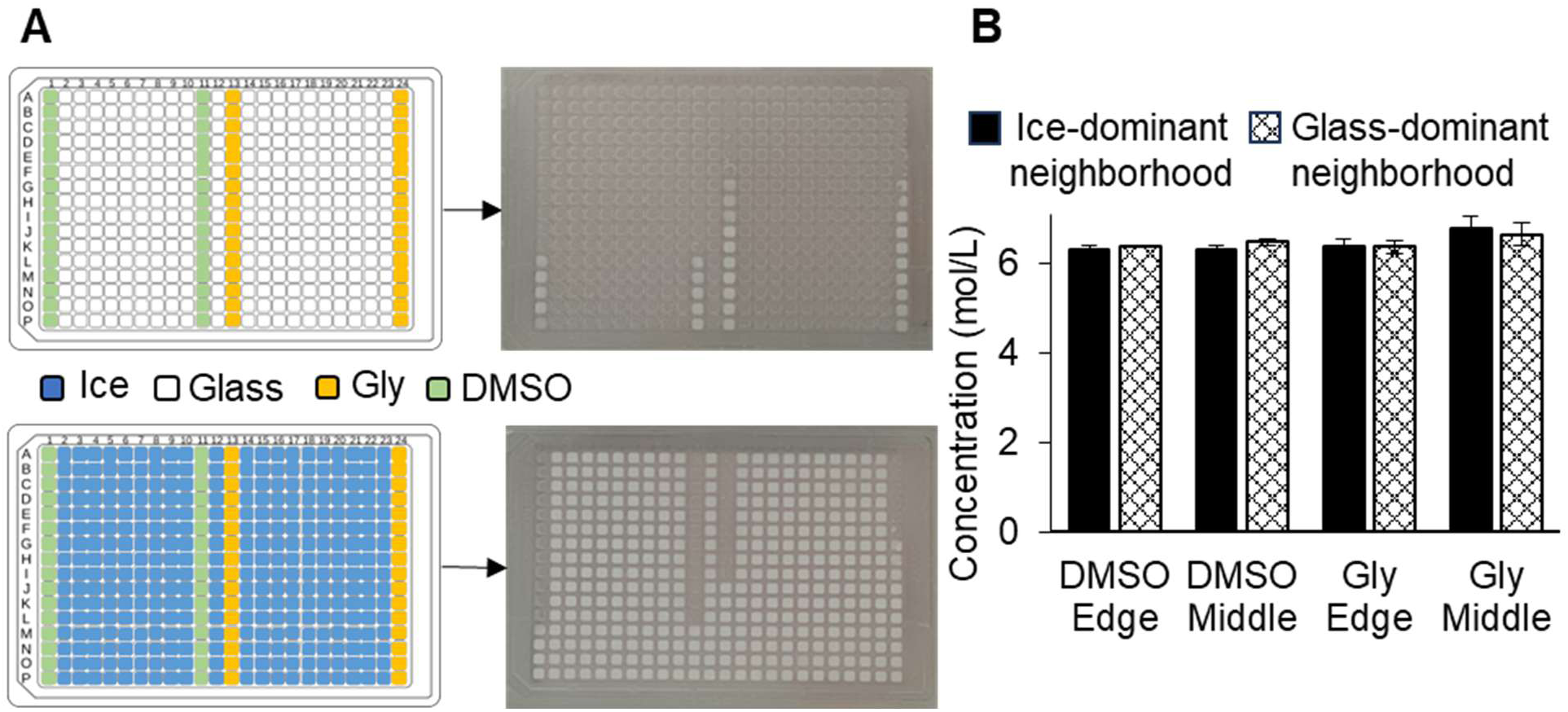
Effects of plate position and local neighborhood. (A) Schematic of the positional/neighborhood validation experiment and representative plate image showing the placement of DMSO and Gly at edge and central locations under different neighborhood conditions. For the DMSO and Gly columns, the concentration varied from 40% w/v at the bottom of the plate (which corresponds to 5.1 mol/L for DMSO and 4.3 mol/L for Gly) to 70% w/v at the top (which corresponds to 9.0 mol/L for DMSO and 7.6 mol/L for Gly). (B) Cv outcomes for DMSO and Gly as a function of plate position and neighborhood state (ice-dominant vs glass-dominant) for plates sealed with a silicone mat, demonstrating that neither parameter measurably alters vitrification behavior under the conditions used (p > 0.05).

### 3.4 Reproducing classical Cv values from 15-mL tube experiments

To benchmark the performance of our platform against established Cv measurements, we compared Fahy’s classical sealed-tube results^22^ to Cv values obtained in our lab using a similar 15-mL tube setup (Figure S6 and Figure S7) and the 384-well automated workflow. Twelve CPA compositions previously tested by Fahy were selected for evaluation (Table S1). Figure 5 shows results for 10 of these CPAs. Two CPAs, BD and AM, were excluded from the plotted comparison because they did not vitrify within the concentration range tested. AM showed only partial signs of vitrification at 56% w/v in our tube experiment and 54% w/v in the 384-well format, whereas BD did not vitrify under either condition.

**Figure 5.**
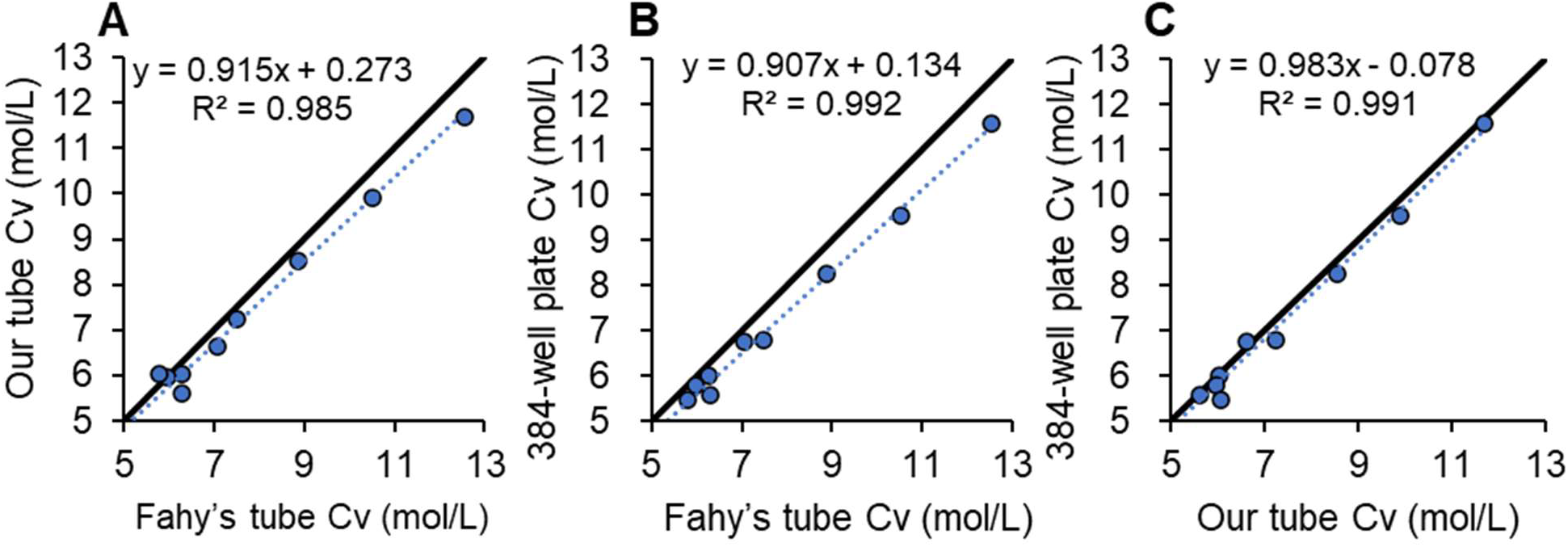
Benchmarking Cv measurements across platforms. (A) Comparison of Cv values measured using Fahy’s sealed-tube method and the 15-mL tube configuration used in this study. (B) Comparison of Cv values measured using Fahy’s sealed-tube method and the 384-well plate automated platform. (C) Comparison of Cv values measured using our 15-mL tube configuration and the 384-well plate automated platform, demonstrating strong agreement between methods. Dotted lines show best-fit lines and the solid lines are identity lines (y = x).

Using the tube-based configuration, we obtained Cv values that closely matched the historical dataset (R² = 0.985). The 384-well plate format also showed strong agreement with the Fahy values (R² = 0.992), and the two methods demonstrated excellent internal consistency (R² = 0.991). These results confirm that the high-throughput assay faithfully reproduces vitrification behavior obtained using classical low-throughput protocols.

### 3.5 Environmental boundary conditions shift Cv values

Because ice can nucleate at air-liquid interfaces,^36^ we hypothesized that the overlying gas environment would affect Cv. To quantify these effects, we evaluated Cv outcomes for 11 CPA compositions under four experimental configurations, as illustrated in Figure 6. Clear differences were observed across configurations for all 11 CPA compositions (Figure S8). In general, plates sealed with silicone mats produced the lowest Cv, while open plates exposed to ambient air produced the highest. For example, the Cv value for PG was 5.47 mol/L under the silicone-mat condition but increased to 7.13 mol/L in the open-top condition. Purging with dry argon in the headspace above open plates resulted in lower Cv values than the standard open-top condition. These data suggest that water vapor in the overlying gas promotes ice formation, possibly by nucleating ice at the gas-liquid interface.

**Figure 6.**
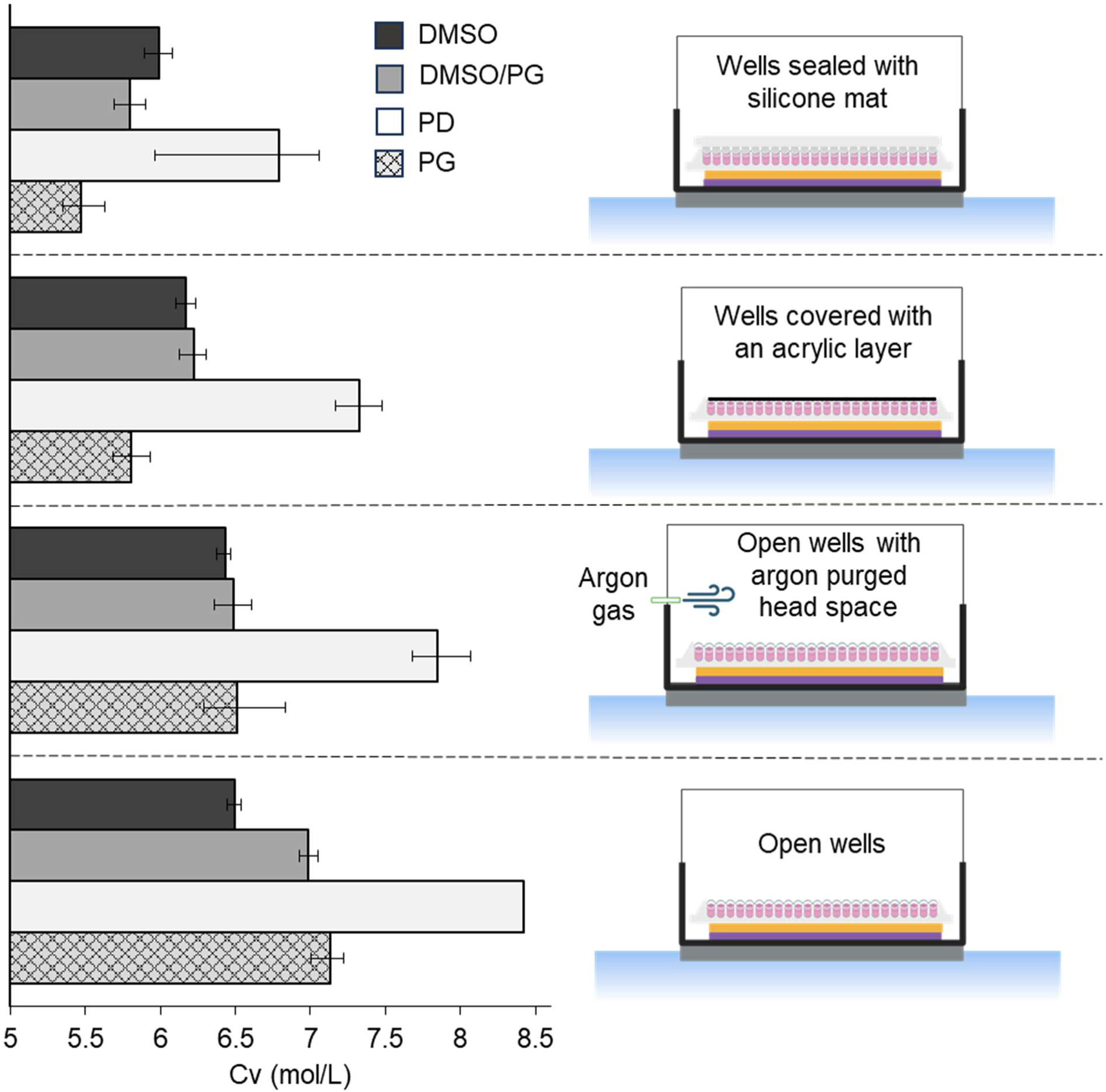
Environmental boundary conditions systematically shift Cv. Cv values were measured under four conditions with different levels of exposure to water vapor in the gas phase above the wells. Error bars show 95% confidence intervals.

To more broadly evaluate the effects of environmental conditions, we tested ∼200 CPA compositions using two configurations (sealed plates and open plates), yielding ∼400 total Cv measurements. These two methods were chosen to illustrate the range of variability introduced by environmental exposure, with the silicone-mat condition representing a controlled and sealed environment, and the open-top condition representing an unsealed, ambient environment. As shown in Figure 7, nearly all compositions for the open-top condition exhibited Cv values that were higher than the silicone-mat condition. Despite this shift, the two methods remained strongly correlated (R² = 0.883), demonstrating that both configurations capture the same underlying trends while differing in absolute Cv values.

**Figure 7.**
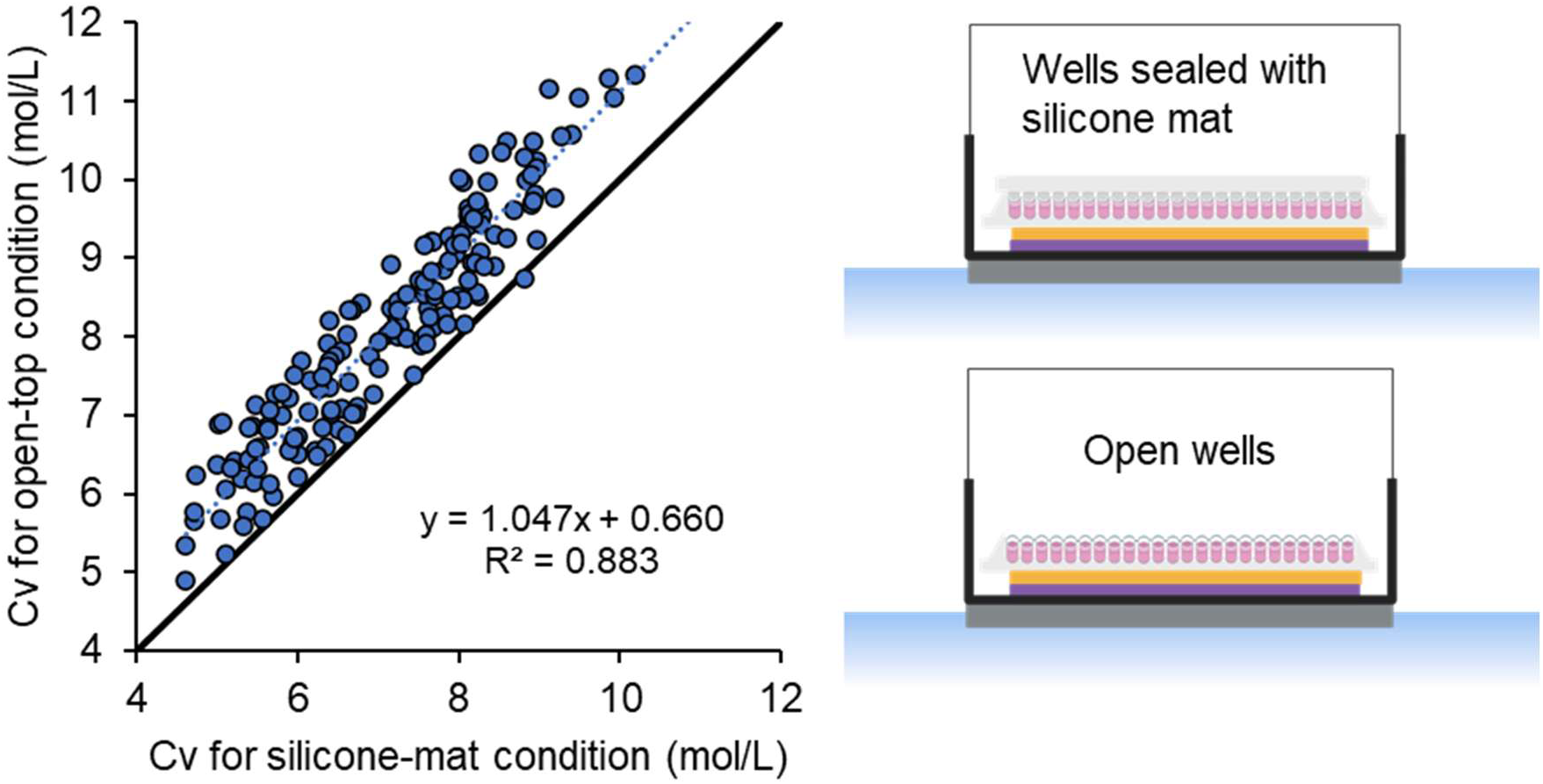
Effect of environmental conditions on Cv across ∼200 CPA compositions. The dotted line is the best-fit line and the solid line is the identity line (y = x).

### 3.6 Relationship between Cv and average molecular weight

Figure 8 examines the effect of molecular weight on Cv. When Cv values were expressed in molar units (mol/L), a clear trend emerged in which Cv decreased systematically with increasing molecular weight (Figure 8A). Overall, Cv decreased by more than a factor of two over the range of molecular weights studied. This indicates that for the CPAs studied here ice suppression *per molecule* is higher for larger molecules, which enables vitrification at a lower molar concentration. In contrast, when Cv was expressed as mass concentration (% w/v), only a slight downward trend with molecular weight was observed. Overall, the data reflects approximately uniform scatter around the average Cv value of 53.34 % w/v (Figure 8B).

**Figure 8.**
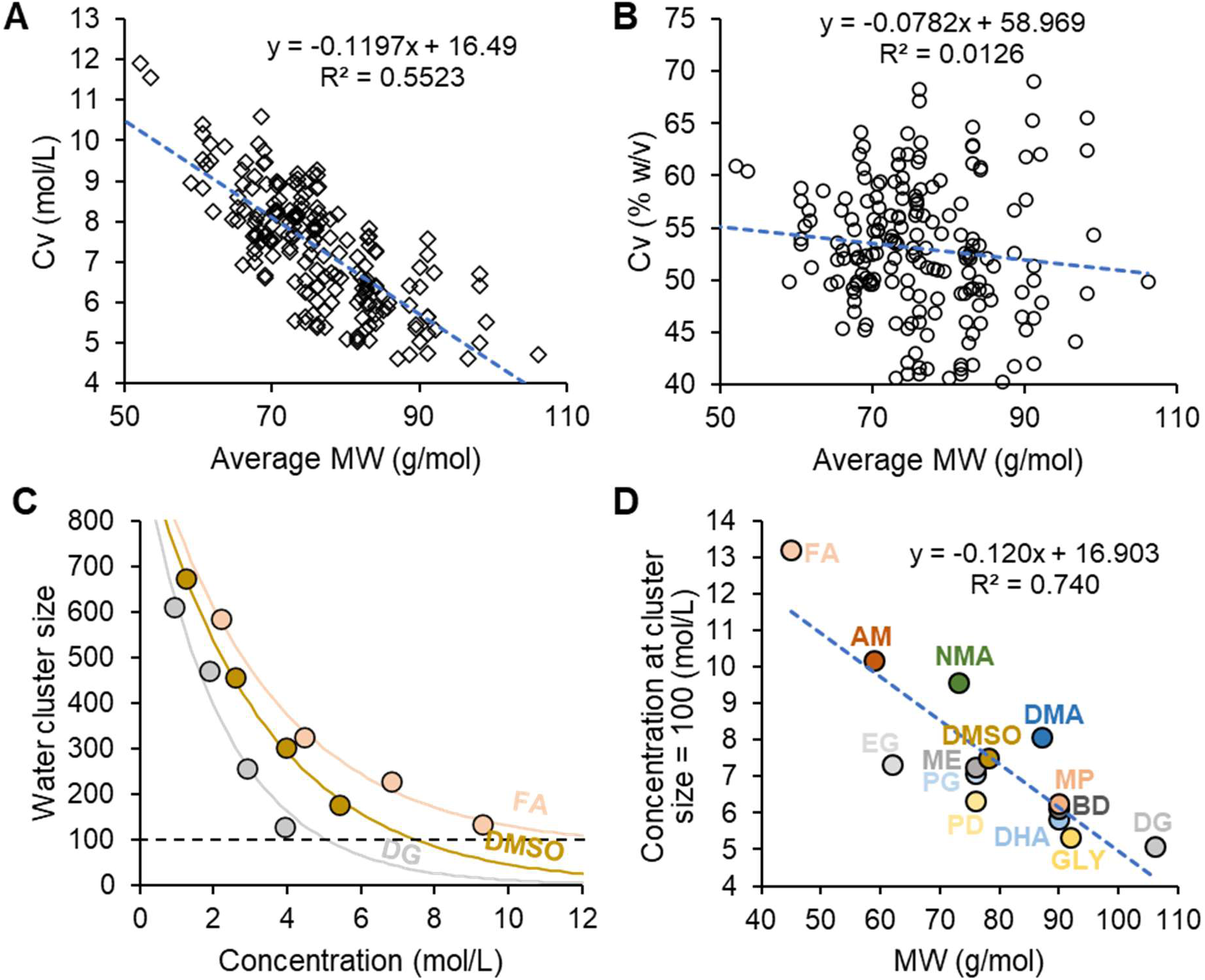
Effect of molecular weight on Cv. (A) Cv in molar units (mol/L) with best-fit line (slope ≠ 0, p = 1.2 × 10^−34^). (B) Cv in mass concentration units (% w/v) with best-fit line (slope ≠ 0, p = 0.12). Cv values are for 384 well plates with silicone sealing mats. (C) Water cluster sizes as a function of concentration for three representative CPAs computed from MD simulations. (D) Effect of molecular weight on the CPA concentration required to achieve a cluster size of 100 water molecules, with best-fit line (slope ≠ 0, p = 7.8 × 10^−5^).

To explore the molecular basis for the observed trends, we performed molecular dynamics simulations to assess the effects of CPAs on the water hydrogen bonding (H-bond) network. As a water H-bond descriptor, we compute the average water cluster size. This was computed from the H-bond connectivity matrix by extracting the participation ratio.^31^ This value shows the number of water molecules in a given cluster. As shown in Figure 8C, CPAs disrupt the water network, resulting in an average water cluster size that decreases as CPA concentration increases. Figure 8D examines the concentration required to achieve a threshold cluster size of 100 water molecules for the 14 CPAs studied here. This value is arbitrary, as other cluster sizes also correlate with Cv, as previously described.^31^ There is a clear downward trend with molecular weight, indicating that larger molecules tend to disrupt water clustering more than smaller molecules. This is consistent with the trend observed for Cv. Together, these results suggest that the enhanced ability of larger CPAs to fragment water clusters may contribute to their lower Cv values by suppressing the cooperative water organization required for ice formation.

### 3.7 Anomalous results in single CPA solutions

Figure 9 illustrates anomalous results for two CPAs (ME and AM) that are consistent with the formation of non-ice crystalline phases in single CPA solutions. For ME, the Cv value of the single-CPA solution is substantially higher than that of its two-CPA mixture with Gly, indicating that the mixture is better at suppressing crystallization. This cannot be attributed to improved suppression of crystallization by Gly, since Gly on its own had a higher Cv than the ME-Gly mixture. This anomalous result is consistent with the known tendency of ME solutions to form a hydrate.^37^ Similarly, AM exhibits behavior consistent with non-ice crystallization. Partial vitrification was observed at ∼9.1 mol/L, but all wells exhibited crystallization below and above this concentration. This suggests two separate modes of crystallization. Together these results suggest that Cv values measured for single-CPA solutions may not reflect the intrinsic ice suppression abilities of CPAs because of the confounding effects of non-ice crystallization.

**Figure 9.**
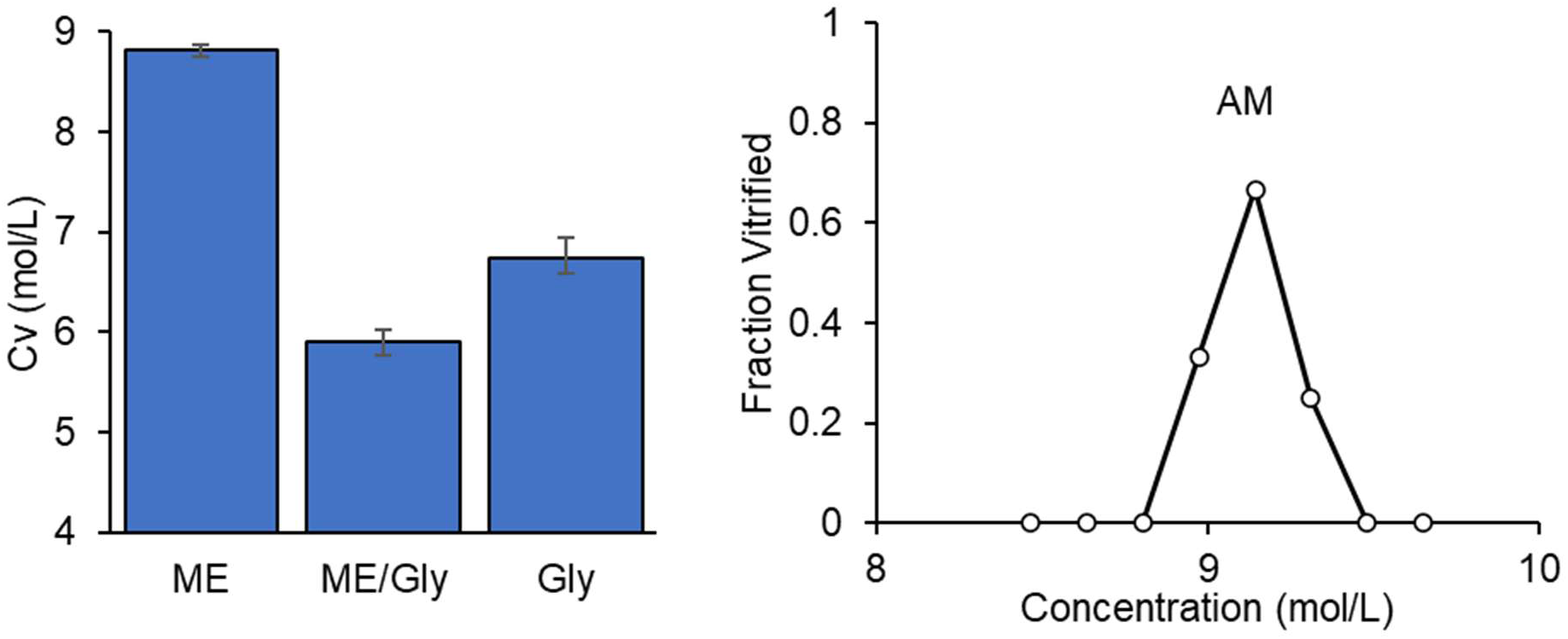
Anomalous results in single CPA solutions suggest the formation of non-ice crystalline phases. Results are for 384 well plates with silicone sealing mats. Error bars show 95% confidence intervals.

### 3.8 Predicting Cv in CPA mixtures

A simple mixture model was used to predict Cv in multi-CPA solutions based on the intrinsic ice suppression abilities of each CPA (Eq. 3). Because of the confounding effects of non-ice crystallization in some single-CPA solutions, only Cv data for solutions containing two CPAs were used to fit the model and estimate the values of the ice suppression parameter *α*. These best-fit *α* values were then used to predict Cv for all the CPA formulations evaluated in this study, including mixtures containing more than two CPAs.

The model demonstrated excellent predictive capability across multiple levels of complexity. For the two-CPA mixtures used to infer the intrinsic ice suppression parameters, predicted and experimentally measured Cv values showed strong agreement (R² = 0.981; Figure 10A). When applied to multicomponent mixtures that were not included during model fitting, the model retained high predictive accuracy, yielding an R² of 0.928 (Figure 10B).

**Figure 10.**
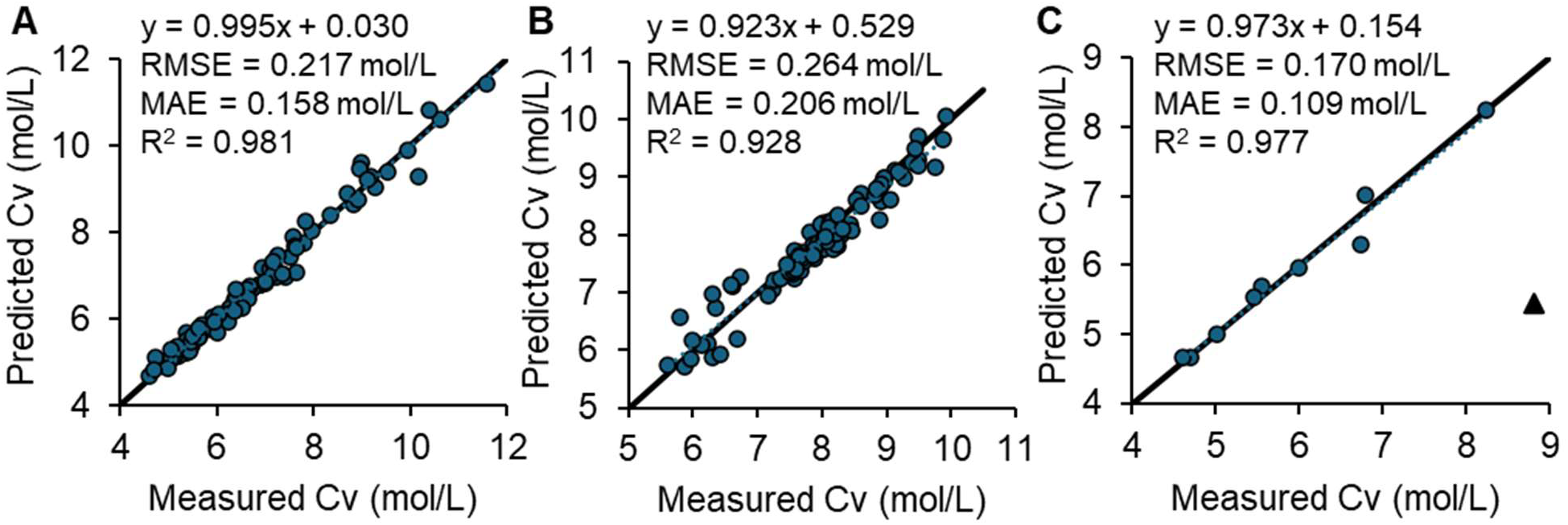
Prediction of Cv using the mixture model (Eq. 3). (A) Predicted versus measured Cv values for CPA mixtures with two CPAs, which were used to fit the mixture-model parameters. (B) Predicted versus measured Cv values for mixtures with more than two-CPAs that were not included in model fitting, demonstrating predictive performance on unseen formulations. (C) Comparison of predicted and measured Cv values for single-CPA solutions. ME is shown with black triangle and excluded from the regression because it is known to form a hydrate.^37^ Cv values are for 384 well plates with silicone sealing mats. See Figure S9 and Table S2 for open-well results.

In addition, comparison of predicted versus experimentally measured Cv values for single CPAs showed strong overall agreement for most of the CPAs (R² = 0.977; Figure 10C). For single-CPA solutions the ice suppression parameter *α* is equivalent to the predicted Cv. Therefore, predicted and measured Cv values are expected to match when ice is the only crystalline phase that forms during cooling. ME is shown separately and was excluded from the regression analysis, as single-CPA solutions are known to form hydrates, which is expected to elevate the measured Cv and obscure ME’s intrinsic ice-suppression behavior. Also excluded are CPAs that did not meet the threshold for vitrification over the range of CPA concentrations tested: AM, BD, DHA, and FA. Table 2 shows the best-fit *α* values for all the CPAs, ranging from 4.67 mol/L for DG and DMA to 15.99 mol/L for FA.

**Table 2.** Best-fit ice suppression parameters.

| CPA | Ice Suppression Parameter, $\alpha$ (mol L<sup>-1</sup>) |
| --- | --- |
| AM | 9.09 |
| BD | 10.51 |
| DG | 4.67 |
| DHA | 8.63 |
| DMA | 4.67 |
| DMSO | 5.97 |
| EG | 8.23 |
| FA | 15.99 |
| GLY | 6.30 |
| ME | 5.44 |
| MP | 5.01 |
| NMA | 5.69 |
| PD | 7.01 |
| PG | 5.53 |

Together, these results demonstrate that the mixture model robustly captures the dominant physicochemical contributions governing ice suppression and accurately predicts vitrification behavior across single-component, two-component, and more complex multi-component CPA formulations.

### 3.9 Evaluating published CPA toxicity data using Cv predictions

Traditional approaches for evaluating CPA toxicity typically assess all formulations at a single, fixed concentration (e.g., testing all CPAs at 6 mol/L).^15, 19^ However, this practice implicitly assumes that the same concentration is appropriate for every CPA, even though different compounds require substantially different concentrations to achieve a glassy state. As a result, measuring toxicity at only one concentration can yield misleading comparisons: some CPAs may be tested far below their vitrification-relevant range, whereas others may be tested far above it.

To determine how closely the concentrations used in prior toxicity studies approached the vitrification concentrations (Cv) determined in this work, we applied our vitrification model to the viability datasets from Ahmadkhani et al.^15^ and Jaskiewicz et al.^19^ The model was used as a reference to estimate Cv for each CPA mixture and to compare these predicted values with the molarities tested in the toxicity assays.

In Figure 11A, viability is plotted against the ratio of the CPA molarity used in the toxicity assay to the predicted Cv for each composition. Ratios near 1 indicate that the toxicity assays were performed at concentrations close to the vitrification threshold, whereas ratios >1 represent cases where viability was assessed at concentrations exceeding the predicted requirement for vitrification. There is a clear trend of decreasing viability as the CPA concentration approaches the vitrification threshold.

**Figure 11.**
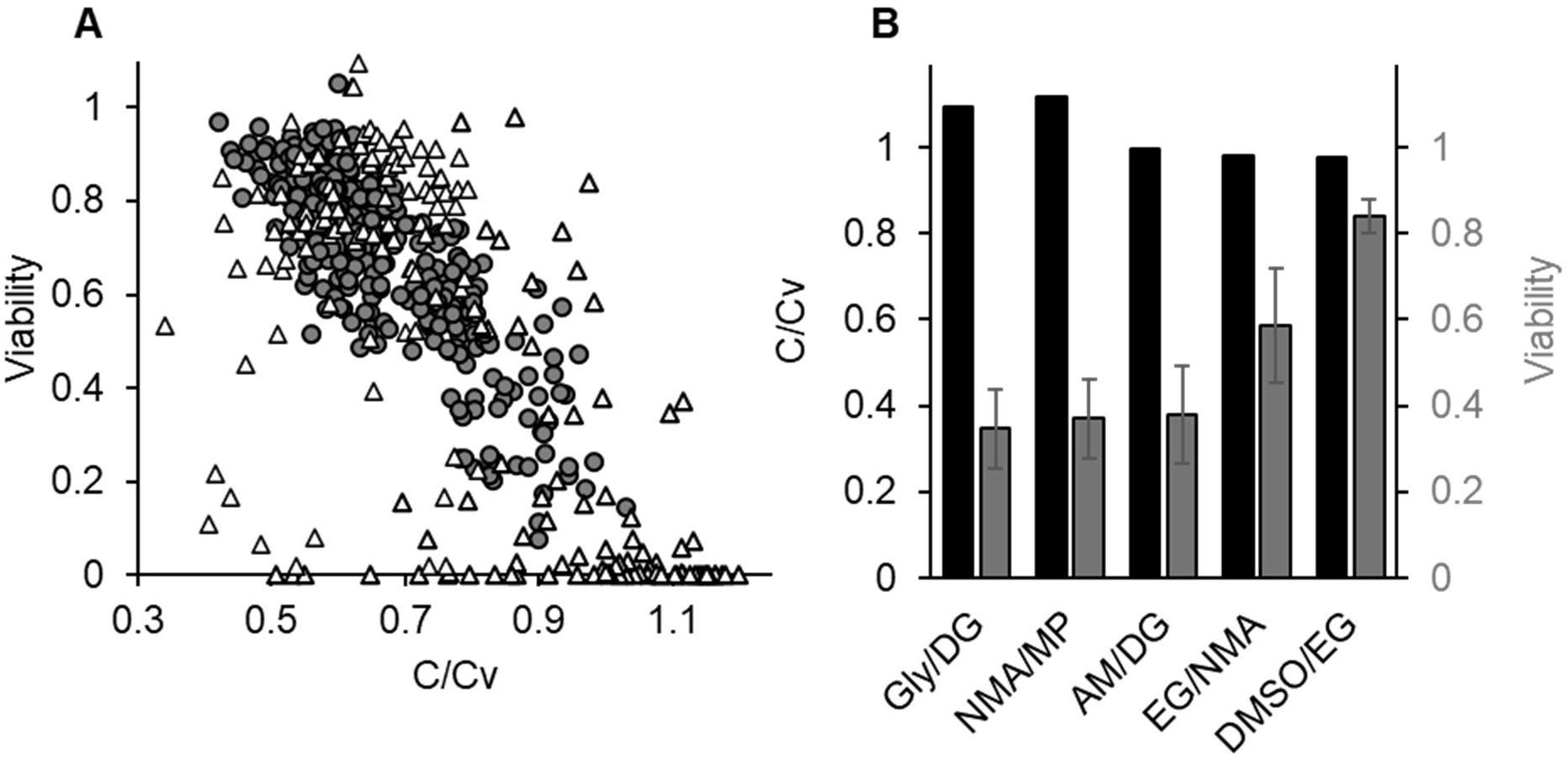
Evaluation of published CPA toxicity data using Cv predictions. (A) Viability after CPA exposure plotted as a function of the concentration ratio C/Cv. Gray circles indicate data adapted from Jackiewicz et al. (320 data points),^19^ and white triangles indicate data adapted from Ahmadkhani et al. (197 data points).^15^ (B) Representative CPA mixtures highlighting formulations tested near or above the predicted vitrification threshold. Black bars represent the concentration ratio C/Cv, and gray bars represent measured viability.

Figure 11B highlights representative mixtures where toxicity measurements were conducted at or above the predicted Cv, as well as compositions that exhibited high viability despite being tested slightly below Cv. For example, Gly/DG exhibited a C/Cv ratio of ∼1.09 with a viability of 0.35; NMA/MP had a ratio of ∼1.12 with a viability of ∼0.37; AM/DG showed a ratio of ∼0.995 with a viability of ∼0.38; EG/NMA had a ratio of ∼0.982 with a viability of ∼0.58; and DMSO/EG exhibited a ratio of ∼0.975 with a viability of ∼0.84.

## 4 Discussion

Stability against ice formation is a critical factor in the design of CPA formulations for vitrification-based cryopreservation. In this study, we report the first high throughput approach for quantifying the CPA concentration required for vitrification (Cv), increasing the speed at which CPA formulations can be evaluated by ∼50x. Using this approach, we made ∼400 Cv measurements based on ∼26,000 individual data points. This far exceeds the ∼45 Cv values in the published literature.^22^ The new method reported here complements recently reported high throughput approaches for assessing CPA permeability and toxicity,^15–19^ filling a key gap in the evaluation of candidate CPA formulations for stability against ice formation. Together, these methods create a platform for discovering new CPAs that balance stability, permeability, and toxicity.

The ability to measure Cv at increased scale creates new opportunities to interrogate the molecular determinants of CPA solution stability. In this study, Cv exhibited a strong dependence on molecular weight, suggesting that larger molecules more effectively suppress ice formation on a per-molecule basis. However, molecular weight alone does not fully account for vitrification behavior. This is illustrated by the comparison between PG and PD, which share the same molecular weight yet display markedly different ice suppression capabilities. These differences point to the influence of additional molecular properties, such as geometry and the distribution of polar functional groups. As larger and more diverse datasets become available, there is substantial opportunity to apply computational chemistry, molecular dynamics simulations, and machine learning approaches to elucidate structure–stability relationships.^38–41^ Such efforts could advance mechanistic understanding of ice formation in aqueous mixtures and ultimately enable prediction of Cv directly from molecular structure.

This study presents a simple model for predicting Cv for CPA mixtures based on the ice suppression properties of each constituent CPA. To estimate the ice suppression parameters for the 14 CPAs evaluated in this study, we fit the model to Cv data for all binary combinations of the CPAs. The resulting fits generalized well, accurately predicting Cv for other mixtures, including formulations with up to seven CPAs. However, a limitation of the model is that it assumes vitrification behavior is governed primarily by suppression of ice formation and therefore does not account for alternative crystallization pathways. This limitation is most evident for ME, which is known to form a hydrate^37^ and consequently deviates from the model predictions (Figure 10C). Moreover, the applicability of the model outside the 14 CPAs considered here has not yet been established.

Despite the promising performance of the Cv model, the fitting procedure scales quadratically with the number of CPAs, which presents a challenge for broader screening of candidate CPA compounds. To address this limitation, we reanalyzed the data using a reduced fitting strategy in which binary combinations with a common set of five CPAs (DMSO, EG, PG, FA, and Gly) were used to estimate ice suppression parameters. The resulting α values differed by less than 3% from those obtained using the full binary dataset, indicating that exhaustive testing of all binary combinations is not required. This reduced approach enables linear scaling of the fitting process, as each new CPA need only be tested in binary mixtures with the common reference set to reliably estimate its ice suppression properties.

The ability to predict Cv for any mixture given the ice suppression parameters of the constituent CPAs has the potential to streamline identification of CPA formulations that are both vitrifiable and nontoxic. As an initial demonstration of this, we re-evaluated published toxicity data in the context of the predicted Cv (Figure 11), revealing some promising compositions with relatively low toxicity near Cv. However, prior CPA toxicity studies have mainly compared CPAs at the same molar concentration,^15, 16, 19^ resulting in varying levels of vitrifiability for different CPAs. Moving forward, the best strategy for optimizing CPA formulation is to evaluate toxicity at Cv.^42^ The high throughput Cv method presented here and the associated Cv model will make it possible to do this at a scale that was previously unattainable.

Our results suggest that formulations containing two or more CPAs are less likely to form non-ice crystalline phases than solutions containing a single CPA. This observation supports the common practice of using multi-CPA cocktails for vitrification. ME provides a clear example of this behavior. In single CPA solutions, ME vitrified only at relatively high concentrations, which is consistent with its known tendency to form a hydrate.^37^ However, when ME was mixed with other CPAs, vitrification occurred at much lower concentrations, suggesting that the presence of additional CPAs disrupted hydrate formation. A similar effect was observed for AM, which exhibited partial vitrification only in a narrow concentration range when used alone but became more stable when incorporated into multi-CPA mixtures. Three additional CPAs (BD, DHA, and FA) failed to vitrify over the entire range of concentrations tested, possibly due to non-ice crystallization. In particular, BD is known to form a hydrate in its meso isomeric form,^43^ which may explain why BD solutions did not vitrify. As shown in Figure S10, we also observed possible non-ice crystallization in two-CPA mixtures containing BD. Given that ice suppression parameters were derived from two-CPA mixtures, we examined whether BD’s anomalous behavior may have introduced bias into the estimated parameters by refitting the model to a reduced dataset excluding BD mixtures. This yielded best-fit ice suppression parameters for the remaining CPAs that were within 2% of the original values (Table S3), confirming the accuracy of our parameter estimates.

The data presented here show that vitrification behavior depends strongly on boundary conditions, with open plates exposed to ambient air consistently requiring higher Cv relative to plates sealed with silicone sealing mats. This mirrors prior observations where “exposed” preparations promoted earlier, more variable ice nucleation, while aseptic, washed conditions delayed nucleation and reduced variability.^44^ In this context, open-top plates represent an exposed, nucleator-rich environment, whereas silicone mat sealing behaves like a cleaner, nucleator-limited environment. A key contributor is likely ice formation at the air–water interface: vapor in the headspace can condense and freeze as frost on cooled surfaces, triggering bulk freezing. Powell-Palm and colleagues have emphasized the role of interfacial and environmental exposure in destabilizing supercooled systems and shown that removing air–water interfaces markedly enhances supercooling stability under isochoric conditions.^36^ Together, these findings demonstrate that environmental exposure and sealing conditions directly influence ice formation behavior and, consequently, the apparent Cv.

Overall, our Cv measurements are consistent with expected trends from previous studies. The Cv values reported by Fahy et al.^20, 22^ correlated strongly with those obtained in the current study. The major exception was BD, which exhibited substantially different Cv behavior. This discrepancy is likely explained by isomeric composition: meso-2,3-butanediol has been reported to form a hydrate and is considered a poor glass former, whereas our BD sample was a mixture with no specification of isomeric proportions.^45, 46^ Variation in isomer content therefore plausibly accounts for the observed differences.

Although the method presented here achieves substantially higher throughput than existing approaches, practical throughput remains constrained to approximately 200 compositions per day, with 12 replicates per composition. Under a five-iteration binary search protocol, this corresponds to roughly 200 Cv measurements per week. The main bottleneck is the time required for automated liquid handling to prepare the CPA mixtures in well plates. It may be possible to increase throughput by further streamlining and optimizing the liquid handling process. Alternatively, throughput could be increased by using more than one liquid handler. Reducing the number of replicates per condition is another potential strategy to increase throughput. For example, decreasing from 12 to 6 replicates would approximately double throughput, at the cost of a ∼40% increase in the standard error of the estimated fraction of vitrified wells; this trade-off may be acceptable in large-scale screening contexts. Additional gains in throughput could be achieved through full automation of image analysis. In the present study, variations in illumination during image acquisition introduced classification inconsistencies that necessitated manual review to verify automated results. Future work should therefore focus on improved control of imaging conditions and the development of more robust image processing approaches to better distinguish vitrified from ice-containing wells, thereby reducing manual intervention and further increasing throughput.

## 5 Conclusion

This work establishes a scalable approach for characterizing vitrification behavior across a broad range of CPA formulations, addressing a key bottleneck hindering development of improved formulations for cryopreservation of complex systems like human organs. The new method yields Cv measurements that align with prior reports while greatly expanding the accessible composition space. Application of this platform yielded ∼400 Cv measurements based on ∼26,000 individual data points, providing insight into the factors affecting vitrification behavior. Vitrification outcomes were strongly influenced by external boundary conditions, with sealed plates consistently favoring glass formation at lower concentrations compared to open plates exposed to ambient air. Analysis across mixtures comprised of 14 CPAs further demonstrated a clear relationship between molecular weight and vitrification efficacy, supporting the view that ice suppression scales with molecular size rather than concentration alone. MD simulations show that water cluster size also depends on CPA molecular weight, providing a molecular-level interpretation of the results. To characterize Cv in mixtures, we developed a model for predicting Cv based on the ice suppression properties of each constituent CPA. This model yielded accurate results for a wide range of CPA formulations, including CPA solutions containing up to 7 CPAs. Using these predicted Cv values, we analyzed 514 literature data points reported for CPA toxicity and identified CPA mixtures that operate near their vitrification threshold while maintaining relatively low toxicity. Collectively, these advances support a more quantitative and scalable approach to CPA discovery and formulation design, with potential to accelerate the development of improved cryopreservation strategies.

## 6 Supporting Information

- Additional methodological details and supplementary results on cooling of 384 well plates, binary search strategy, image analysis, positional and neighborhood effects in 384 well plates, cooling of 15 mL tubes, compositions for benchmarking Cv measurement, environmental boundary condition effects, and Cv prediction for open plates.
- Experimental data, including CPA mixture compositions, measured Cv values for each composition, and the fraction of wells that vitrified for each composition

## 7 Acknowledgements

This work was supported by funding from the National Science Foundation (EEC 1941543). CRB acknowledges support from the National Institutes of Health (R35GM133359). We are grateful to the following students who contributed to this work: Adam Mayo, Emma Anderson, Tomi Fagbemi, Robby Donnatelli, Maxwell Sisson, Natalie Allen, Guillermo Vazquez Galindo, Morgan Johnson. Several figures include icons or drawings created using BioRender.com: Figure 1, 2, 6, 7, and Figure S3. This is a SAWIAGOS project.

