## Supporting Information for "A high throughput platform for measuring and predicting vitrification behavior in multicomponent aqueous solutions"

### Cooling setup for 384 well plates

The high throughput platform for Cv measurement includes an apparatus for simultaneous cooling of three 384 well plates (Figure S1). The plates rest on stacked acrylic and copper sheets inside a stainless-steel pan, and a cube-shaped acrylic cover encloses the pan to control the ambient environment. This setup is placed in a Styrofoam box and cooled using liquid nitrogen.

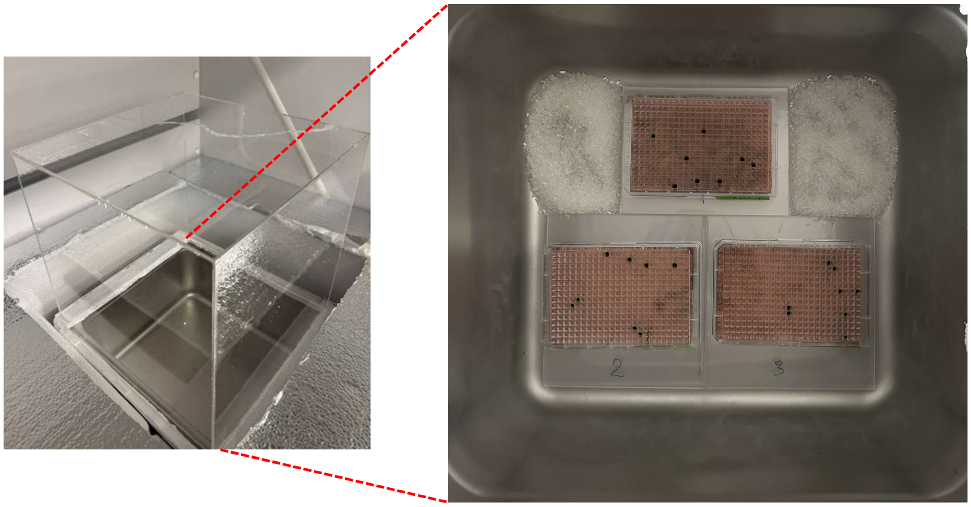

*Figure S1. Images of cooling setup. Left: Full setup showing stainless steel pan with acrylic cover. The pan does not contain the 384-well plates in this image. Right: Top view of stainless steel pan containing three 384-well plates.*

### Cooling rate measurements in 384-well plates

Figure S2 shows cooling-rate measurements obtained from a 384-well plate during Cv experiments. Cooling rates were calculated from the temperature change between approximately

−20 °C and −120 °C for multiple well locations distributed across the plate. Measurements were performed for three plates, and all data points are shown.

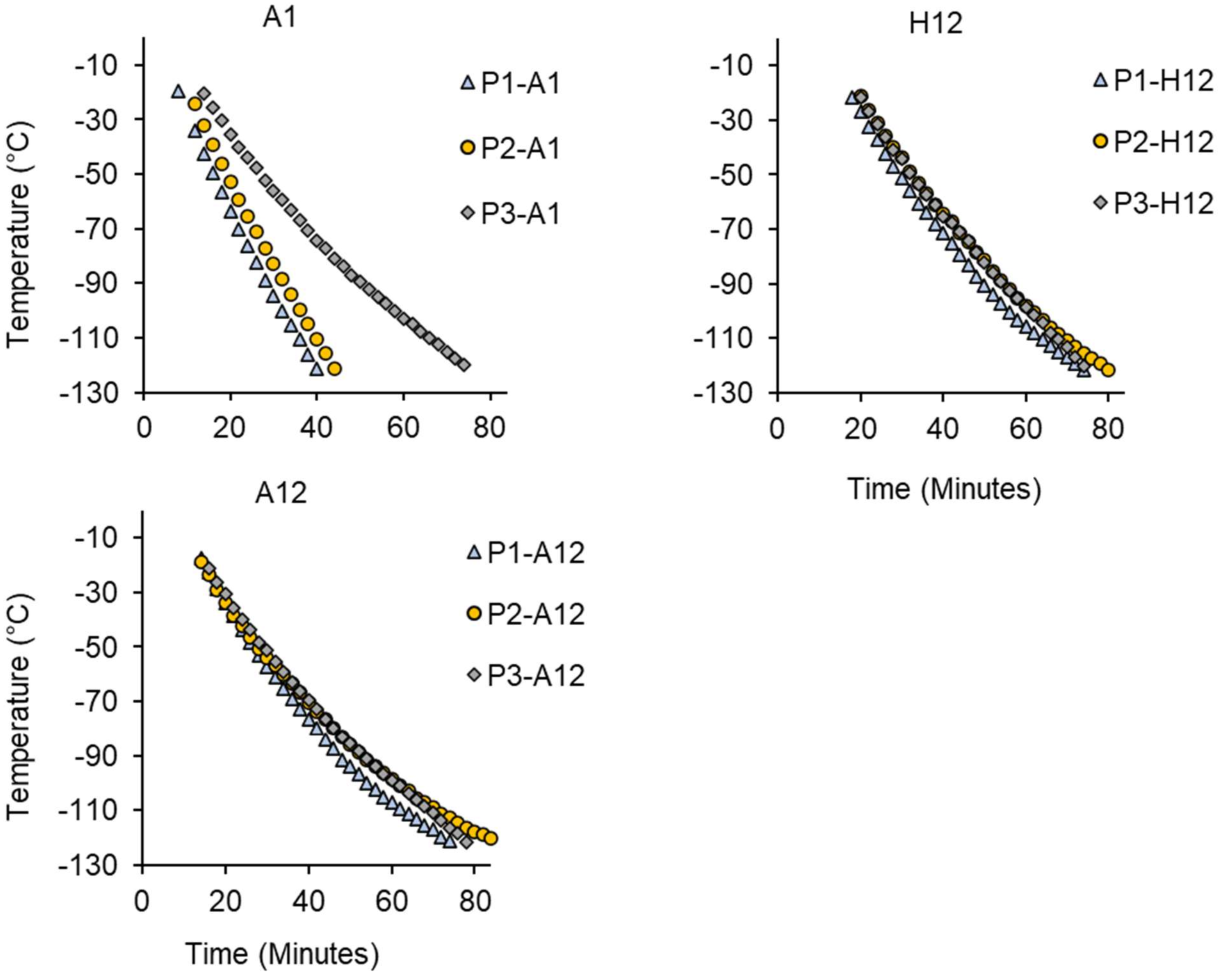


*Figure S2. Cooling-rate measurements across the 384-well plate. Cooling rates calculated from the temperature change between approximately −20 °C and −120 °C at multiple well locations across the plate. Data are shown for three independent replicates (i.e., three plates, P1-P3).*

**Comparison of traditional linear scanning and binary search strategies for Cv determination**

Traditional Cv measurement relies on linear scanning, in which a series of concentrations are tested sequentially (e.g., 38-70% w/v in 1% increments). This approach can require over 30 individual measurements to determine Cv. To streamline Cv measurement, we implemented a binary search strategy that iteratively narrows the concentration range based on vitrification outcome (Figure S3). At each step, the midpoint concentration is tested, and the next test is chosen based on whether the outcome is ice or glass. For example, starting from an initial test at 54% w/v (i.e., the midpoint between 38% and 70% w/v), an ice outcome leads to testing 62% w/v, which is the midpoint between 54% and the upper bound of 70% w/v. If instead the outcome is a glass, this leads to testing 42% w/v, the midpoint between 54% and the lower bound of 38% w/v. Repeating this process five times resolves Cv to 1% precision. This is more than 6x lower than the 33 measurements required for a full linear scan.

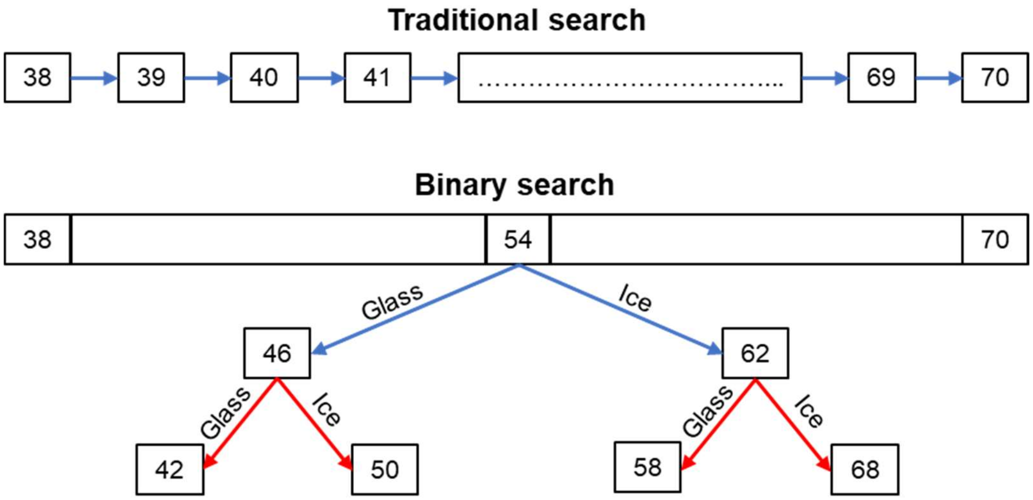


*Figure S3. Binary-search strategy for efficient Cv determination. Schematic comparison of traditional linear scanning and a binary-search approach to find Cv in the range 38% to 70% w/v. A full linear scan requires 33 measurements to achieve 1% resolution, while only 5 iterations of the binary search are required to achieve this resolution.*

**Image analysis**

Images of 384 well plates were analyzed using the process illustrated in Figure S4. To identify the location of each well, the user is asked to click in the center of reference wells at the top left, top right, bottom left, and bottom right. This allows creation of a grid of well centers for the entire plate. Circular regions of interest are defined around these grid centers (Figure S4B), and wells are automatically classified as either frozen or vitrified based on analysis of the average and standard deviation of the intensity in the region of interest. An image is then displayed showing wells classified as frozen using a red dot (Figure S4C). The user is then asked to manually correct the automated well classifications by clicking on the well to toggle on/off the red dot, resulting in a manually corrected image (Figure S4D). The resulting well classifications are saved as a csv for downstream analysis.

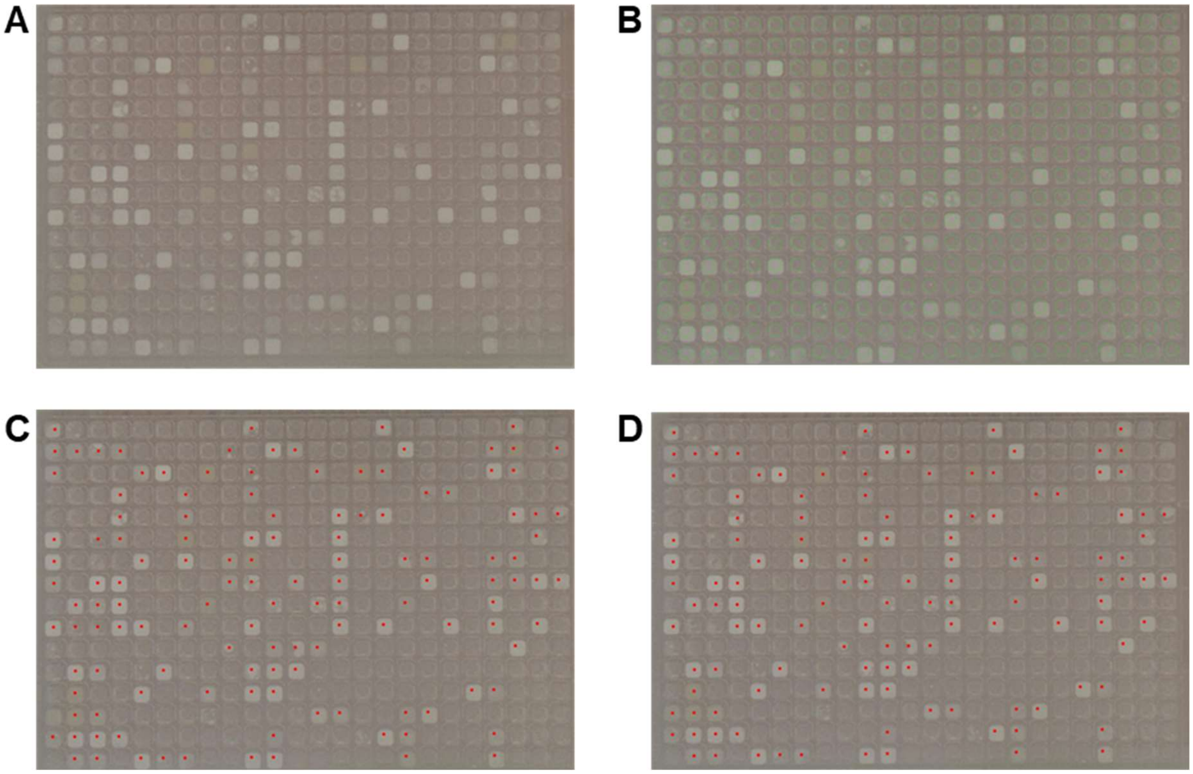


*Figure S4. Semi-automated process for analysis of well plate images. (A) Original cropped image of well plate. (B) After the user identifies the location of four reference wells in the corners of the plate, circular regions of interest are defined for each well. (C) Automated analysis of pixel intensity in these regions is used to classify each well as frozen (red dot) or vitrified. (D) The user then manually corrects the automated classification by clicking on misclassified wells, resulting in a final corrected image.*

**Supplementary validation of positional and neighborhood effects using an acrylic cover**

To complement the silicone-mat experiments presented in the main text, we performed a positional and neighborhood validation experiment under a closed-top acrylic cover configuration. DMSO and glycerol (Gly) were dispensed at both central and edge well locations to assess whether plate position or local neighborhood influenced vitrification outcomes under this alternative covered-plate condition. Neighborhoods were classified as ice-dominant or glass-dominant based on whether the majority of surrounding wells exhibited ice formation or vitrification, respectively. Across all tested conditions, no detectable differences in vitrification outcomes were observed for either CPA as a function of plate position or neighborhood classification (Figure S5, $p > 0.05$). These results indicate that, similar to the silicone-mat configuration, the acrylic-covered condition does not introduce measurable positional or neighborhood bias in Cv determination, further supporting the robustness of the 384-well platform.

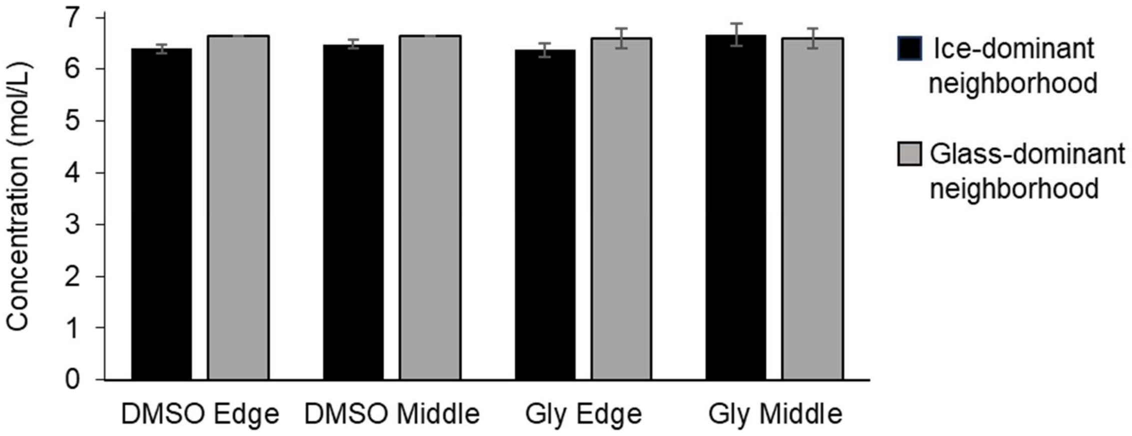


*Figure S5. Vitrification outcomes for DMSO and GLY as a function of plate position and neighborhood state (ice-dominant vs glass-dominant), demonstrating that neither parameter measurably alters vitrification behavior under the conditions used ($p > 0.05$).*

**Cooling setup for tube-based Cv measurements**

Tube-based Cv measurements were performed using a custom setup designed to cool 21 samples simultaneously. CPA test solutions were prepared from 70% w/v stock solutions to generate the desired concentration series. Each solution was loaded into a 15-mL conical tube, which was then inserted into a 50-mL conical tube to control the cooling rate.

As shown in Figure S6, a circular opening matching the outer diameter of the 15-mL tube was created in the cap of the 50-mL tube. This allowed the inner tube to be suspended securely while preventing the 15-mL cap from passing through the opening. The resulting geometry ensured that the 15-mL tube did not contact the walls of the 50-mL tube, thereby minimizing variability in heat transfer and providing consistent thermal insulation across samples. The use of the outer 50-mL tube reduced the effective cooling rate relative to direct immersion, enabling controlled assessment of vitrification behavior.

Following cooling, the 15-mL tubes were removed from the outer tubes and immediately recorded on video to document vitrification outcomes. Representative still images were extracted from the recorded videos for each formulation and used for visual classification. Samples were categorized as vitrified (glass) or ice-forming based on the presence of bulk ice. The appearance of a thin ice layer confined to the upper ~1 mL of the solution was not used as a criterion for ice classification, as this feature was observed in certain CPAs due to interfacial effects rather than bulk crystallization.

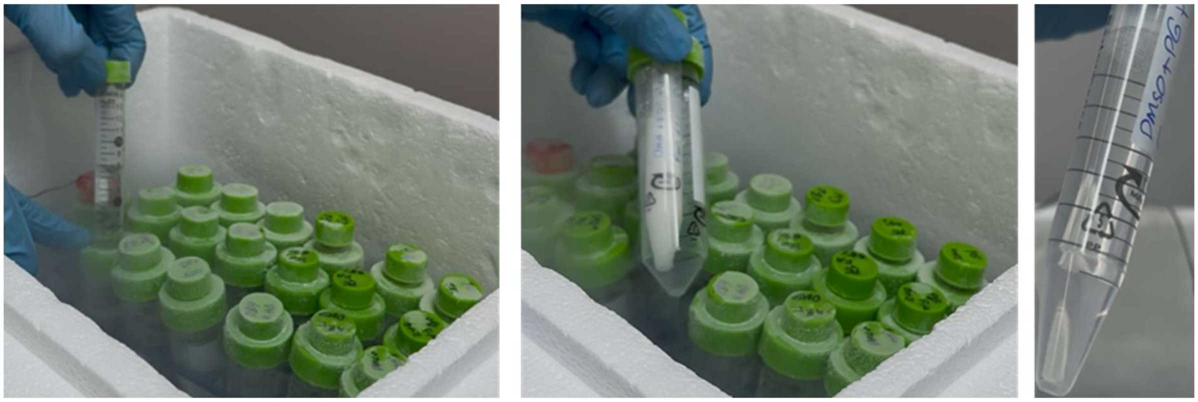

*Figure S6. Tube-based cooling setup for Cv measurements. The 15-mL tube is suspended within a 50-mL tube to reduce cooling rate and minimize thermal variability. Post-cooling samples are recorded and visually classified as vitrified or ice-forming based on bulk ice formation.*

**Cooling rate in tube-based Cv experiments**

To measure cooling rate, a small hole was drilled in the cap of each 15-mL tube, allowing insertion of the thermocouple probe into the solution. Cooling rates were recorded at multiple positions within the tube to assess consistency of temperature profiles across the sample volume. As shown in Figure S7, cooling rates measured at multiple vertical positions within the 15-mL tube revealed a pronounced spatial dependence. The highest cooling rate was consistently observed at the lower conical tip, whereas cooling rates measured along the cylindrical body of the tube were comparatively similar to one another. This behavior is expected due to the increased surface-to-volume ratio at the conical tip, which enhances heat transfer during cooling. Cooling rates measured along the cylindrical region of the tube were used to represent the effective cooling rate for tube-based Cv measurements reported in this study.

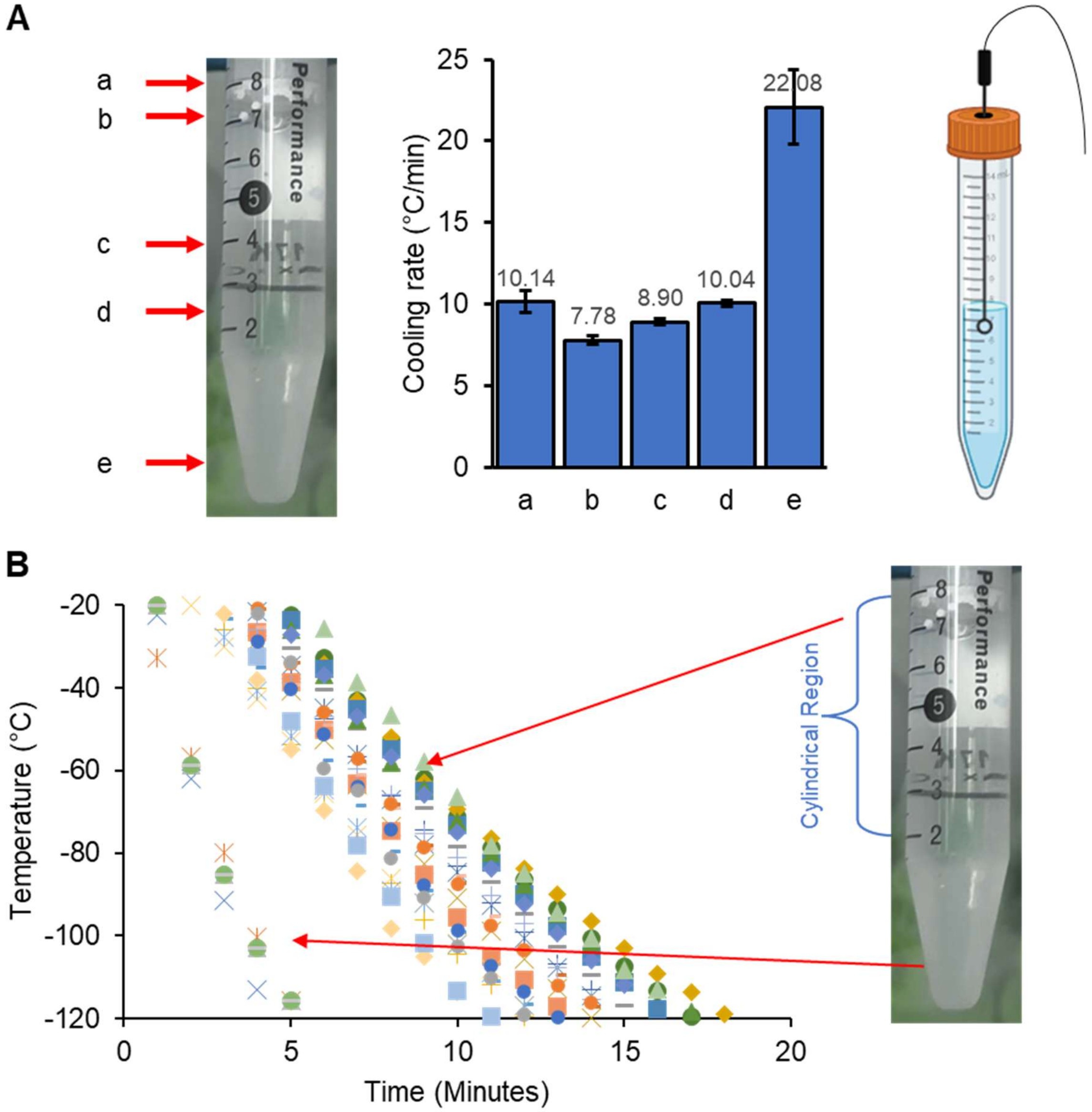


*Figure S7. Spatial cooling-rate measurements in the tube-based Cv setup. (A) Representative image of a 15-mL tube after cooling, with red arrows indicating the vertical positions at which cooling rates were measured using an inserted thermocouple. (B) Cooling rates measured at the indicated positions (−20 °C to −120 °C), showing substantially higher cooling at the lower conical tip and similar cooling rates along the cylindrical region of the tube, which was used to define the representative cooling rate for tube-based Cv measurements.*

### CPAs selected from Fahy's tube experiments for benchmarking our methods

Twelve CPA compositions previously tested by Fahy [1] were used for comparison of his tube-based Cv measurements to those obtained using our tube platform and our 384 well platform (Table 2). Eleven of these CPA compositions were also used to systematically investigate the effects of environmental conditions on Cv measured in the 384-well plate format (BD was omitted from this portion of the study because it did not vitrify in tube experiments at concentrations up to 70% w/v). We examined four different environmental boundary conditions: plates sealed with silicon mats, plates covered with an acrylic layer, open plates exposed to ambient air, and open plates with the head space purged by dry argon. For each CPA, multiple concentrations (in increments of 1% w/v) were prepared and dispensed into randomized well locations using our high-throughput workflow. After completion of this experiment, the plates sealed with a silicone mat were reused to examine the effect of cooling rate in the 384-well plate format.

*Table S1. CPA compositions selected from Fahy's tube experiments [1] to benchmark our methods, and the concentration ranges used to assess their vitrification behavior. Of these 12 compositions, the first 11 were used to systematically investigate the effects of environmental conditions on Cv in 384 well plates.*

| | **Concentration range (% w/v)** | | | | |
|---|---|---|---|---|---|
| **CPA** | Wells sealed with silicone mat | Wells covered with an acrylic layer | Open wells with argon purged head space | Open wells | 15 mL tubes |
| PD | 51-58 | 51-60 | 53-60 | 53-70 | 53-58 |
| DMSO | 45-52 | 45-52 | 45-52 | 45-52 | 46-49 |
| DMSO/PG* | 42-50 | 42-52 | 44-52 | 44-70 | 44-46, 50 |
| DMA | 38-45 | 38-47 | 40-47 | 40-47 | 38-42, 46 |
| FA/PG* | 54-62 | 54-62 | 54-62 | 54-70 | 52-58, 60 |
| NMA | 36-43 | 36-46 | 39-46 | 39-46 | 39-41, 46 |
| EG | 48-55 | 48-58 | 51-58 | 51-70 | 51-56 |
| PG | 38-47 | 38-49 | 40-49 | 58-70 | 43-46 |
| GLY | 57-66 | 57-66 | 57-66 | 57-70 | 57-65 |
| EG/FA* | 58-65 | 58-65 | 58-65 | 40-70 | 59-63, 66 |
| AM | 50-57 | 50-61 | 54-61 | 54-70 | 52-56, 61 |
| BD | Binary search | n/a | n/a | Binary search | 46, 50, 67, 70 |

* Each CPA was present at an equal mass concentration (% w/v).

#### Environmental boundary conditions shift Cv values

We assessed vitrification behavior for the first 11 compositions shown in Table S1 for four different environmental conditions: plates sealed with silicone mats, plates covered with a layer of acrylic, open plates exposed to ambient air, and open plates with the head space purged by dry argon. The results of these experiments are shown in Figure S8.

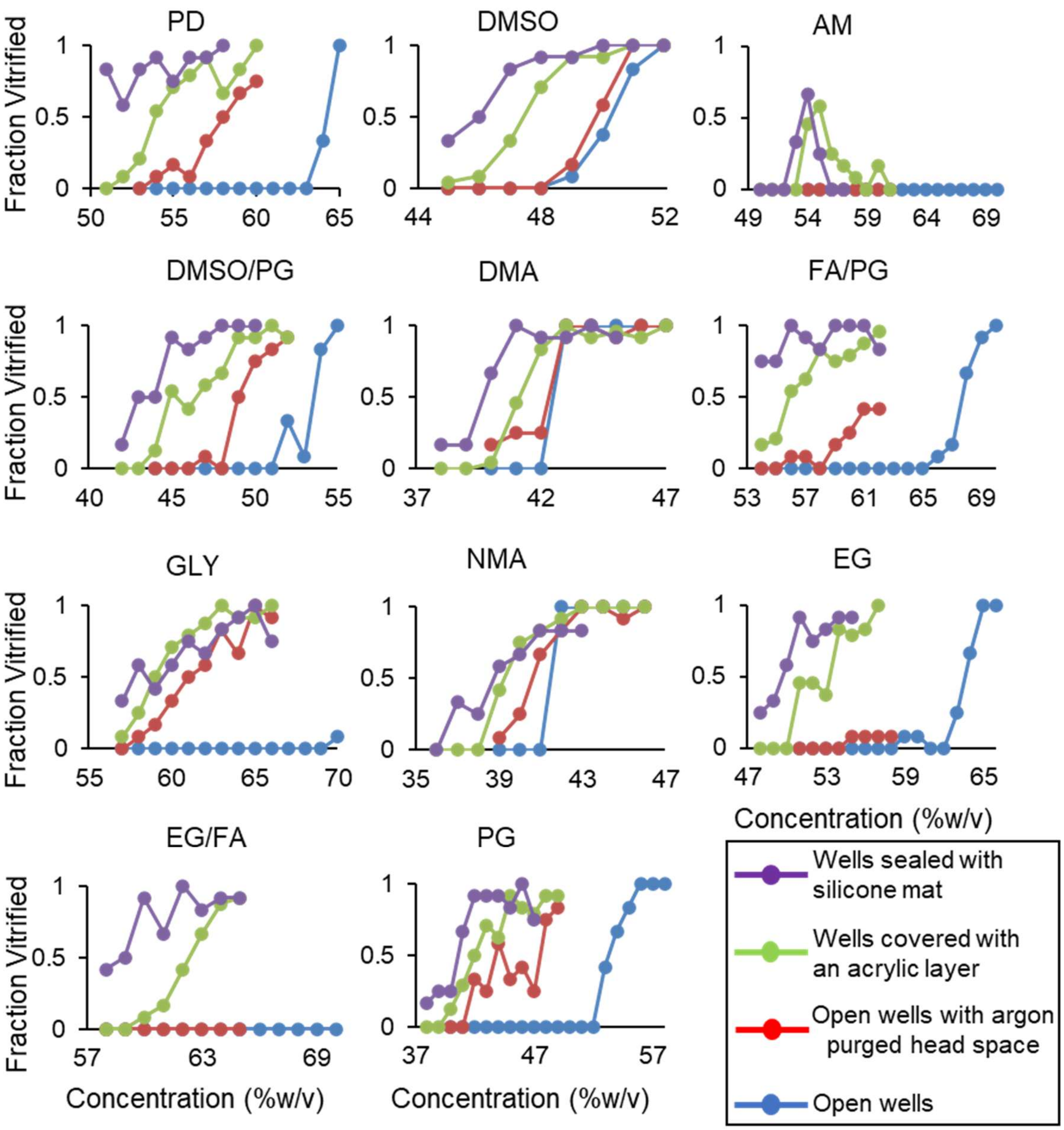


*Figure S8. Effect of environmental conditions on vitrification behavior in 384 well plates, showing a general trend of lower Cv in sealed plates compared to open plates. The compositions in Table S1 were tested, and Cv was determined using a sigmoid fit as described in the main text. BD was also tested for plates sealed with silicone mats and open plates, but BD solutions did not vitrify over the range of concentrations tested for open plates and only partially vitrified at the highest concentration for plates sealed with silicone mats.*

**Prediction of Cv for open plates**

The analysis of the Cv data for open plates was similar to that described in the main manuscript for plates sealed with a silicone mat. To train the model, we used the Cv data for two-CPA mixtures, resulting in the best-fit ice suppression parameters in Table S2. As shown in Figure S9 the best-fit predictions match the data for two-CPA solutions, with a coefficient of determination ($R^2$) of ~0.98. However, the agreement between predictions and Cv data for mixtures with more than two CPAs is relatively poor ($R^2$ = 0.82).

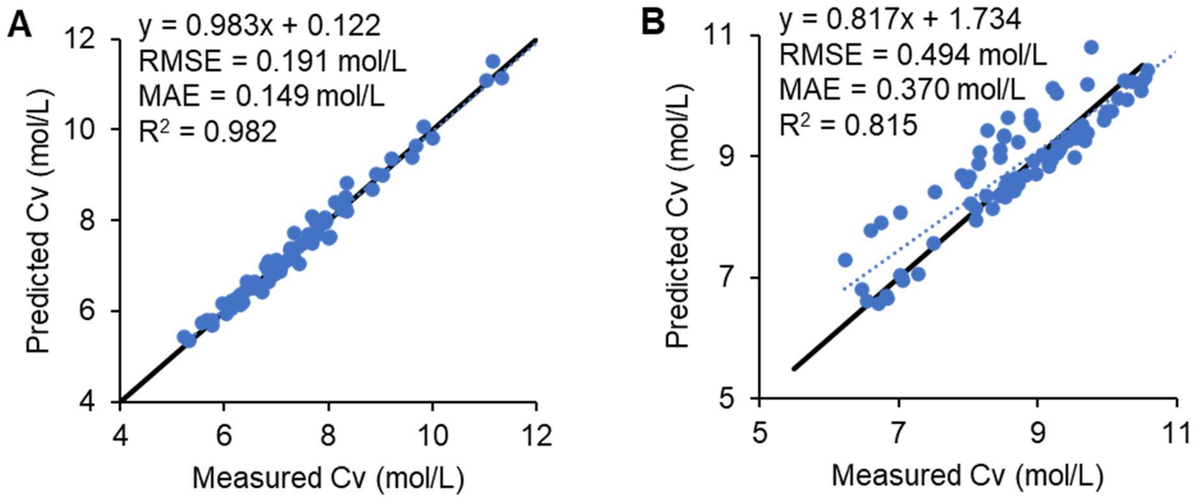


*Figure S9. Prediction of Cv using the mixture model (Eq. 3) for open plates. (A) Predictions for two-CPA solutions, which were used to train the model. (B) Predictions for multi-CPA solutions with more than two CPAs. Black lines are identity lines and dotted lines are best-fit lines.*

*Table S2. Best-fit ice suppression parameters for open plates.*

| CPA | Ice Suppression Parameter, α (mol $L^{-1}$) |
|---|---|
| AM | 9.44 |
| BD | 12.42 |
| DG | 5.77 |
| DHA | 9.33 |
| DMA | 5.05 |
| DMSO | 6.57 |
| EG | 10.24 |
| FA | 18.35 |
| GLY | 7.68 |
| ME | 6.71 |
| MP | 6.59 |
| NMA | 5.81 |
| PD | 8.46 |
| PG | 7.07 |

**Anomalous results for CPA mixtures containing BD**

To investigate the source of deviations between measured and predicted Cv values, we examined the influence of BD, which is known for the strong hydrate-forming capability of its meso isomer.[2]

All CPA mixtures containing more than two CPAs were analyzed, and mixtures containing BD were compared with those lacking BD under both silicone-mat-covered and open-well conditions.

Under silicone-mat-covered conditions, the largest deviations were observed for mixtures containing three or four CPAs when BD was present (Figure S10A). In contrast, mixtures containing more than four CPAs exhibited reduced deviation, likely due to the lower fractional contribution of BD within the overall mixture.

Under open-well conditions, however, the presence of BD resulted in pronounced deviations from the model across nearly all CPA mixtures containing more than two CPAs (Figure S10B). Notably, exclusion of BD-containing mixtures restored strong agreement between predicted and measured Cv values, with $R^2$ improving from approximately 0.82 to 0.97.

In both cases, Cv values were lower than predicted, which indicates that the ice suppression parameter determined from two-CPA mixtures underestimates BD's ability to suppress ice formation in multi-CPA mixtures with more than two CPAs. This is consistent with non-ice crystallization (e.g., hydrate formation) in two-CPA solutions containing BD and concomitant elevation of BD's apparent ice suppression parameter.

To determine whether BD's anomalous behavior biased the ice-suppression parameters of other CPAs, model fitting was repeated excluding mixtures containing BD for both open-well and silicone-mat-covered configurations. In all cases, the resulting changes in fitted ice-suppression parameters were less than 2% (Table S3). These results support the overall accuracy and robustness of the predictive framework presented in this study.

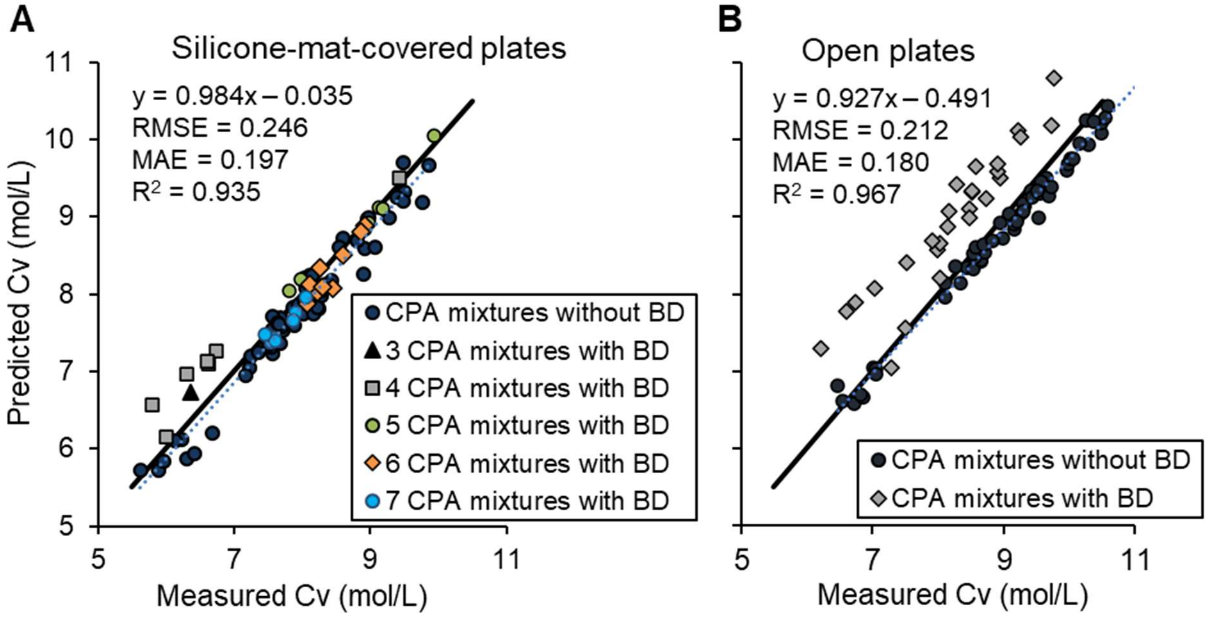


*Figure S10. Effect of BD on model predictions for mixtures with three or more CPAs. (A) Silicone-mat–covered plate. (B) Open-well plate. In both cases, the trend line (dotted line) was fitted using only CPA mixtures that did not contain BD. The solid line shows the identity line.*

*Table S3. Effect of BD on best-fit ice suppression parameters (α, mol $L^{-1}$) for plates sealed with silicone mats and for open plates. Fits were performed for the entire set of binary CPA combinations, or for the subset excluding CPA mixtures containing BD.*

| CPA | Silicone mat | Silicon mat, excluding BD | Open | Open, excluding BD |
|---|---|---|---|---|
| AM | 9.09 | 9.19 | 9.44 | 9.44 |
| BD | 10.51 | | 12.42 | |
| DG | 4.67 | 4.67 | 5.77 | 5.77 |
| DHA | 8.63 | 8.63 | 9.33 | 9.33 |
| DMA | 4.67 | 4.68 | 5.05 | 5.05 |
| DMSO | 5.97 | 6.00 | 6.57 | 6.60 |
| EG | 8.23 | 8.20 | 10.24 | 10.24 |
| FA | 15.99 | 15.99 | 18.35 | 18.35 |
| GLY | 6.30 | 6.35 | 7.68 | 7.68 |
| ME | 5.44 | 5.39 | 6.71 | 6.71 |
| MP | 5.01 | 5.00 | 6.59 | 6.59 |
| NMA | 5.69 | 5.70 | 5.81 | 5.77 |
| PD | 7.01 | 7.07 | 8.46 | 8.46 |
| PG | 5.53 | 5.47 | 7.07 | 7.07 |
